# BET Inhibition Enhances TNF Mediated Anti-Tumor Immunity

**DOI:** 10.1101/2021.02.15.429851

**Authors:** Lisa C. Wellinger, Simon J. Hogg, Dane M. Newman, Thomas Friess, Daniela Geiss, Jessica Michie, Kelly M. Ramsbottom, Marina Bacac, Tanja Fauti, Daniel Marbach, Laura Jarassier, Phillip Thienger, Axel Paehler, Leonie A. Cluse, Conor J. Kearney, Stephin J. Vervoort, Jane Oliaro, Jake Shortt, Astrid Ruefli-Brasse, Daniel Rohle, Ricky W. Johnstone

## Abstract

Targeting chromatin binding proteins and modifying enzymes can concomitantly affect tumor cell proliferation and survival, as well as enhance anti-tumor immunity and augment cancer immunotherapies. By screening a small molecule library of epigenetics-based therapeutics, BET bromodomain inhibitors (BETi) were identified as agents that promote the anti-tumor activity of CD8^+^ T-cells. BETi sensitized diverse tumor types to the cytotoxic effects of the pro-inflammatory cytokine TNF. By preventing the recruitment of BRD4 to p65-bound *cis*-regulatory elements, BETi suppressed the induction of inflammatory gene expression, including the key NF-κB target genes *BIRC2* (cIAP1) and *BIRC3* (cIAP2). Disruption of pro-survival NF-κB signaling by BETi led to unrestrained TNF-mediated activation of the extrinsic apoptotic cascade and tumor cell death. Administration of BETi in combination with T-cell bispecific (TCB) antibodies increased bystander killing of tumor cells and enhanced tumor growth inhibition *in vivo* in a TNF-dependent manner. This novel epigenetic mechanism of immunomodulation may guide future use of BETi as adjuvants for immune oncology agents.

**STATEMENT OF SIGNIFICANCE:** Manipulating the epigenome is an evolving strategy to enhance anti-tumor immunity. We demonstrate that BET bromodomain inhibitors potently sensitize solid tumors to CD8^+^ T-cell killing in a TNF-dependent manner. This immunomodulatory mechanism can be therapeutically leveraged to augment immuno-oncology therapies, including TCB antibodies and immune checkpoint blockade.

## INTRODUCTION

Dysregulation of epigenetic processes that control gene expression in normal physiology are causatively implicated in the pathogenesis of cancer and targeting chromatin modifying enzymes and binding proteins has emerged as a therapeutic strategy in oncology (*1, 2*). One recent example are Bromodomain and Extra-Terminal domain (BET) proteins (BRD2, BRD3, BRD4, and BRDT), which bind to acetylated histones and transcription factors (TFs) at the chromatin interface (*3*). BET proteins localize to active *cis*-regulatory elements throughout the genome and functionally tether transcriptional co-activators, such as P-TEFb (*4*) to RNA polymerase II and stimulate gene expression. Small molecule BET inhibitors (BETi) displace BET proteins from the chromatin and lead to selective suppression of transcription. In genetically- and histologically-diverse tumor types, BETi induce potent anti-proliferative and pro-apoptotic effects that are mechanistically linked to suppression of master lineage-specific TFs, such as c-MYC, and their associated transcriptional programs (*5*). Currently, multiple BETi are under clinical development in hematological and solid malignancies, both as single-agents and in combination with other anti-cancer agents. Here we describe the novel, non-covalent pan-BET inhibitor RG6146 (TEN-010, RO6870810), which was derived through optimization of the widely used prototypical BETi, JQ1, and has been under clinical investigation in context of acute myeloid leukemia and myelodysplastic syndrome (NCT02308761), B-cell lymphomas (NCT03255096), ovarian and breast cancers (NCT03292172), and multiple myeloma (NCT03068351).

Effective immunological responses against cancer are dependent on the orchestrated recruitment and activation of distinct subsets of immune cells to the tumor microenvironment (TME). Immune oncology (IO) therapies have been developed to (i) augment or re-engage the endogenous anti-tumor immune response, using monoclonal antibodies or small molecules, or (ii) induce a tumor antigen-specific immune response by genetic engineering of immune cells employed as cellular therapies. Clinical outcomes to specific IO agents have been correlated with infiltration and activity of effector CD8^+^ T-cells within the TME (*6, 7*), highlighting their central role in anti-tumor immunity. Despite compelling activity in subsets of patients, *de novo* (primary) or acquired resistance against immunotherapy ultimately limits the efficacy of IO agents (*8-11*). While resistance mechanisms of tumor cells to CD8^+^ T-cells and IO agents are multi-faceted (*12, 13*), epigenetic mechanisms to silence the expression of genes controlling antigen processing and presentation and tumor antigens are recurrently identified (*14*). In addition to exploiting tumor cell intrinsic dependencies, it is now well established that epigenetic therapies can functionally modulate anti-tumor immune responses (*15*). The anti-tumor activity of the first-generation BETi, JQ1, was shown to be dependent upon the adaptive immune system in syngeneic models of lymphoma (*16*) and ovarian cancer (*17*). In these models, and others (*18*), BET proteins were shown to co-regulate the expression of immune checkpoint ligand, Programed death-ligand 1 (PD-L1), which can be transcriptionally downregulated by BETi. Furthermore, combination therapies incorporating BETi and IO agents, including checkpoint blockade (*16, 19*) and adoptive T-cell transfer (*20*), have highlighted the ability of BETi to augment both endogenous and IO-induced anti-tumor immunity. Whilst it is clear that tumors utilize BET proteins to drive the expression of key immune evasion molecules, such as PD-L1, there are limited functional studies examining how BETi may directly modulate killing by cytotoxic lymphocytes (CTLs).

Here, we have discovered that BETi, such as RG6146 and JQ1, can directly augment the anti-tumor activity of CD8^+^ T-cells independently of perforin-mediated CTL cytolysis. BETi reprogram the transcriptional response of tumor cells to the pro-inflammatory cytokine TNF leading to potent sensitization to TNF-dependent extrinsic apoptosis. Together, this highlights an additional immune-dependent mechanism by which BETi promote anti-tumor responses, further reinforcing the notion that BETi may be rationally partnered with IO agents.

## RESULTS

### Tumor antigen-specific CD8^+^ T-cell killing is enhanced by BET inhibition

To identify chromatin modifying enzymes and binding proteins that could be targeted therapeutically to augment anti-tumor CTL responses, a boutique library of small molecule epigenetic therapies was screened using an *in vitro* tumor cell-CD8^+^ T-cell co-culture system. The screen was performed using human colon adenocarcinoma HCT-116 cells, presenting either an immunodominant NLV or EBV peptide in the context of HLA-A*02:01, which were incubated with individual compounds from the small molecule library in the presence of human healthy donor-derived CMV-specific T-cells for 48 hours (Figure 1A). We assessed differential viability between tumor cells presenting NLV-(specific T-cell antigen) and EBV (non-specific T-cell antigen) peptides to identify molecules capable to enhancing antigen-specific CTL killing. These analyse revealed that among those small molecules tested, multiple BETi (including RG6146, OTX015, Mivebresib, ABBV744, iBET151 and JQ1) were capable of significantly potentiating CTL-dependent tumor killing of HCT-116 cells presenting the NLV peptide (Figure 1B). In these assays, co-culture with BETi increased tumor cell sensitivity to CD8^+^ T-cell killing to the same extent as the SMAC mimetics Birinapant and LCL-161, which have been previously reported to sensitize tumors to CD8^+^ T-cell killing (*21*). Small molecule inhibitors of the P300/CBP bromodomain modules, SGC-CBP30 and I-CBP-112, were not capable of increasing CTL-dependent tumor killing to the same extent, indicating specificity for bromodomain module of the BET sub-family. Whilst Histone Deacetylase inhibitors (HDACi) Entinostat, Panobinostat and Vorinostat exhibited anti-tumor activity as a single-agent (Supplementary Fig 1A) and this effect was further enhanced in the presence of CTL (Fig 1B). Together, these CD8^+^ T-cell co-culture screening assays have revealed that BETi can directly promote antigen-specific CTL-dependent tumor killing.

**Figure 1.**
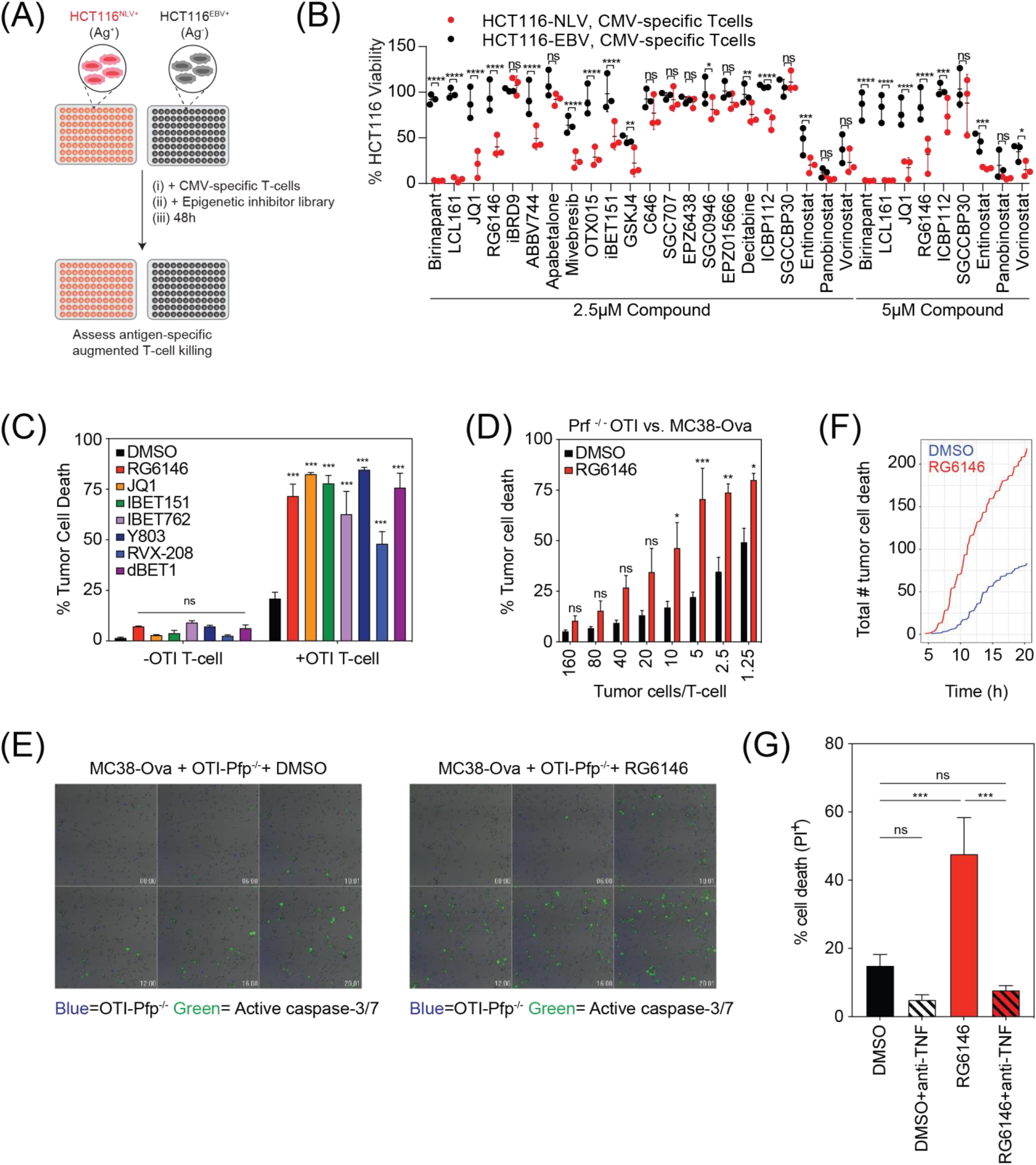
Identification of epigenetic therapies that promotes T-cell mediated anti-tumor immunity. **(A)** Schematic of CD8^+^ T-cell-tumor cell co-culture screen in which NLV or EBV peptide loaded HCT-116 cells were co-cultured in the presence of CMV-specific T-cells and 2.5-5μM of a small molecule library of epigenetic therapies for 48 hours. **(B)** Viability of HCT-116 cells following co-culture with epigenetic therapies or SMAC mimetics and CMV-specific T-cells. Data represents three independent experiments and was normalized to DMSO control. Statistical significance denotes difference between NLV and EBV loaded HCT-116 cells (2-way ANOVA, *p<0.05, **p<0.01, ***p<0.001, ****p<0.0001). **(C)** MC38-Ova cells were co-cultured in 1µM BET inhibitor (10µM for RVX-208) or DMSO in the presence or absence of activated OTI T-cells (20:1 tumor cells/OTI) for 18 hours prior to assessment of tumor cell death by flow cytometry (propidium iodide positivity). Data represents mean of three independent experiments. Statistical significance denotes difference between DMSO and each BET inhibitor (2-way ANOVA, ***p<0.001). **(D)** MC38-Ova cells were co-cultured in 2.5µM RG6146 or DMSO in the presence of increasing ratios of perforin deficient (Prf^-/-^) OTI T-cells for 18 hours prior to assessment of tumor cell death by flow cytometry (propidium iodide positivity). Data represents mean of three independent experiments. Statistical significance denotes difference between DMSO and RG6146 at each ratio (2-way ANOVA, *p<0.05, **p<0.01, ***p<0.001) **(E)** Live cell imaging of MC38-Ova cells seeded in chamber slides in the presence of RG6146 (2.5µM) or DMSO and exposed to Prf^-/-^ OT-I T-cells labelled with CTV. Tumor cell death was visualized with a cleaved caspase-3/7 reporter added to the co-culture media. Representative still images at the indicated time-points are depicted (hr:min). **(F)** Temporal quantification of live cell imaging data of MC38-Ova cells becoming positive for cleaved caspase-3/7 in the presence of Prf^-/-^ OTI T-cells within 20 hours. **(G)** MC38-Ova cells were co-cultured with Prf^-/-^ OTI T-cells (10:1 tumor cells/OTI) for 18 hours in the presence of RG6146 (or DMSO vehicle) and anti-TNF neutralizing antibodies prior to assessment of tumor cell death by flow cytometry (propidium iodide positivity. Data represents mean of three independent experiments (2-way ANOVA, ***p<0.001).

In parallel, the ability of BETi to augment antigen-specific CTL anti-tumor responses was assessed with a small molecule screen in an orthogonal mouse co-culture system that utilized effector CD8^+^ T-cells derived from OTI transgenic mice (referred to as OTI T-cells) that recognize syngeneic (C57BL6-derived) tumor cells expressing the Ovalbumin (Ova) cognate antigen. Recapitulating the findings from the human co-culture system, RG6146, and other structurally distinct BETi, significantly potentiated the CTL-dependent killing of murine colon adenocarcinoma cell line, MC38-Ova (Figure 1C and Supplementary Figure 1B). These results demonstrated that BETi can promote the anti-tumor activity of effector CD8^+^ T-cells independently of species or the specific TCR being utilized. To define the mechanisms by which BETi promoted CTL-dependent killing of tumors, OTI:MC38-Ova co-culture assays were repeated using perforin-deficient (Prf^-/-^) OTI T-cells, which are incapable of direct perforin-dependent cytolysis. Interestingly, BETi were capable of potentiating the anti-tumor activity of Prf^-/-^ OTI T-cells towards MC38-Ova targets (Figure 1D), suggesting a prominent role for secreted cytokines in mediating anti-tumor immunity. This finding was further validated at the single cell level using time-lapse microscopy. CellTrace™ Violet (CTV)-labeled Prf^-/-^ OTI T-cells were co-cultured with MC38-Ova cells in the presence or absence of RG6146, where cleavage and activation of caspase-3/7 was visualized using an exogenous fluorescent reporter. Tumor cell death was increased in the presence of RG6146 compared to vehicle, as visualized in representative still images (Figure 1E), time-lapse videos (Supplementary Videos 1-2), as well as in direct quantification of cell death events across multiple fields (Figure 1F). Collectively, these data demonstrate that BETi promote CTL-driven tumor cell killing independent of the perforin-mediated granule exocytosis pathway.

Following recognition of cognate tumor antigens, CD8^+^ T-cell activation leads to potent acquisition of effector functionality, including production and secretion of pro-inflammatory cytokines. We recently defined the relative contributions of pro-inflammatory cytokines in MC38:OTI co-culture systems, which revealed that TNF, but not IFN-γ, was essential for tumor cell killing (*12*). Consistent with these findings we demonstrated that neutralization of TNF abrogated tumor cell killing by OTI T-cells, but also completely limited the effect of RG6146 in the co-culture assay (Figure 1G). These findings were verified in the co-culture setting with HCT-116 and CMV-specific T-cells (Supplementary Figure 1C). Importantly, BETi treatment was not associated with increased intracellular accumulation of TNF in CD8^+^ T-cells, nor enhanced secretion of TNF into the co-culture supernatant (Supplementary Figure 1D-E). Thus, these data suggested that BETi sensitized tumor cells to the cytotoxic effects of TNF secreted by CD8^+^ T-cells.

### BET inhibition broadly potentiates the cytotoxic activity of TNF

To investigate this possibility, a diverse range of human and mouse tumor cell lines were then screened for the ability of RG6146 to modulate the cytotoxic effects of TNF. TNF potentiated RG6146-mediated tumor cell death in approximately 40% (36/89) of human cell lines tested (Figure 2A). Sensitization to TNF-mediated killing by RG6146 and other BETi was further validated in human HCT-116 and MKN45 (Supplementary Figure 2A), and mouse MC38-Ova and E0771-Ova (Figure 2B-C and Supplementary Figure 2B) solid tumor lines in a time- and dose-dependent manner. Using computational synergy modeling (*22*), we found that combinations of TNF and RG6146 elicited synergistic induction of cell death in MC38-Ova cells (Figure 2D). Finally, the potent pro-apoptotic effects of RG6146 and TNF in combination were evident in patient biopsy-derived colorectal cancer (CRC) specimens propagated *ex vivo* as organoids embedded within matrigel (Figure 2E). Notably, those cancer cell lines and primary patient samples that were limited in their response to the combination of RG6146 and TNF generally exhibited primary resistance towards TNF. For example, two additional CRC patient-derived organoids were only modestly sensitive or refractory to TNF and the combination with RG6146 (Supplementary Figure 2C-D). Similarly, the murine mammary adenocarcinoma (AT3) cells were highly refractory to TNF and showed no significant induction of tumor cell death upon addition of BETi (Supplementary Figure 2B). Thus, the mechanism by which BETi potentiates the pro-apoptotic effects of TNF was ultimately dependent upon an inherent sensitivity to TNF. Nonetheless, in all models tested where inherent sensitivity to TNF was observed, tumor cell death was significantly enhanced by concurrent BET protein inhibition.

**Figure 2.**
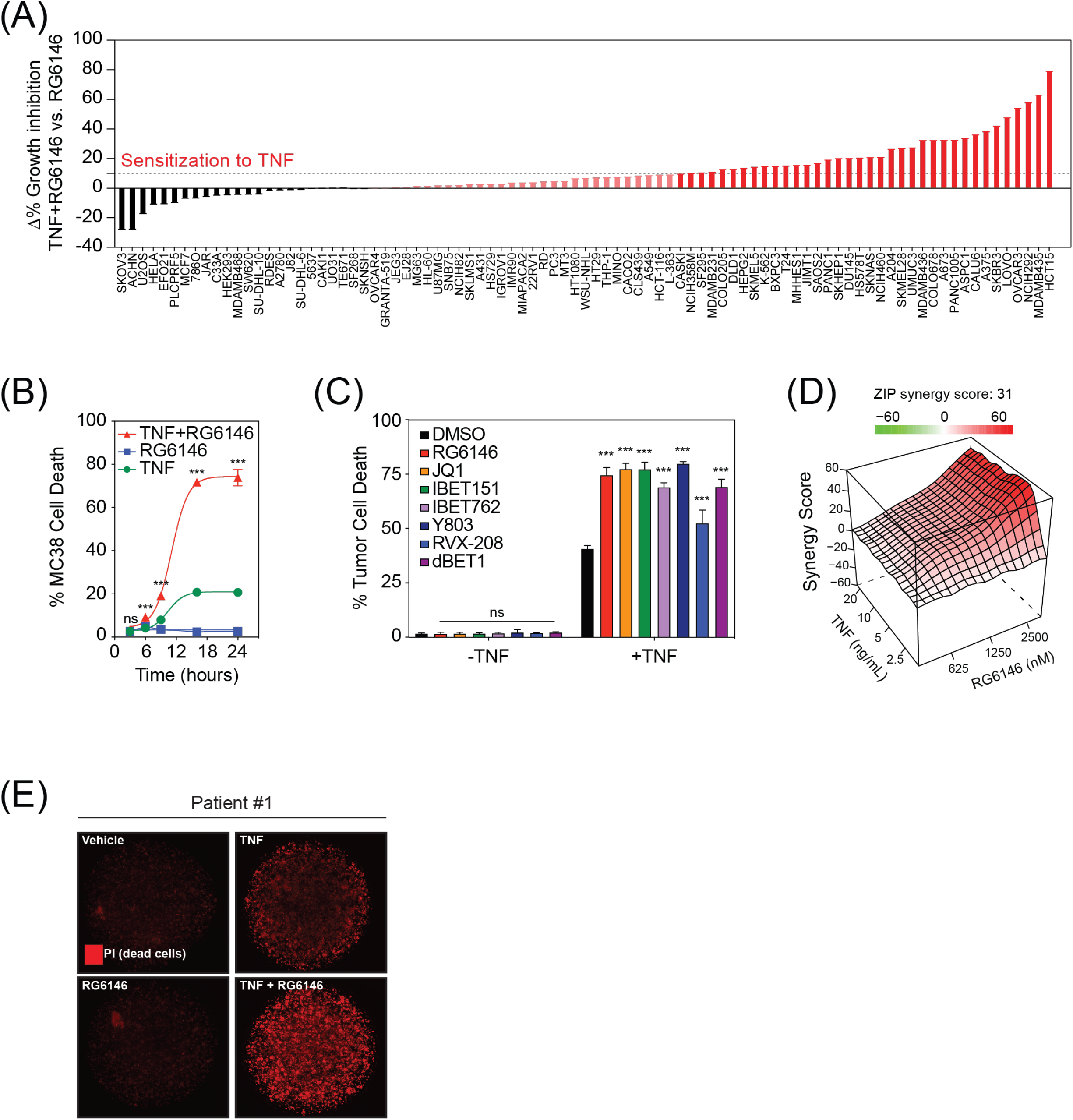
BET proteins functionally protect tumor cells from the cytotoxic effects of TNF. **(A)** Viability of 89 tumor cell lines following treatment with recombinant human TNF (15ng/mL) and RG6146. Maximal growth inhibition (GI_MAX_) of RG6146 was subtracted from GI_MAX_ of RG6146 and TNF combination treatment. Difference in growth inhibition (%) reveals cell lines sensitive (red), unchanged (light red) and resistant (black) to the combination of RG6146 and TNF. **(B)** MC38-Ova cells were cultured in the presence of RG6146 (2.5µM) and recombinant murine TNF (10ng/mL) for indicated time points prior to the assessment of cell death by flow cytometry (propidium iodide positivity). Data represents mean of three technical replicates from representative experiment. Statistical significance denotes difference between TNF and TNF+RG6146 (2-way ANOVA, ***p<0.001). **(C)** MC38-Ova cells were co-cultured in 1µM BET inhibitor (10µM for RVX-208) or DMSO in the presence or absence of recombinant murine TNF (10ng/mL) for 18 hours prior to assessment of tumor cell death by flow cytometry (propidium iodide positivity). Data represents mean of three independent experiments. Statistical significance denotes difference between DMSO and each BET inhibitor (2-way ANOVA, ***p<0.001). **(D)** MC38-Ova cells were cultured in increasing concentrations of RG6146 and recombinant murine TNF for 24 hours prior to assessment of tumor cell death by flow cytometry (propidium iodide positivity). SynergyFinder was utilized to calculate synergy scores using the Zero Interaction Potency (ZIP) reference model. Upward deviation on the y-axis (red) indicates positive synergy across the dose response matrix. (**E**) Fluorescence microscopy of patient-derived colorectal cancer (CRC) organoid embedded in Matrigel and exposed to combinations of RG6146 (2.5µM) or DMSO and recombinant human TNF (15ng/mL) in media containing propidium iodide (PI).

The interaction of TNF with TNF receptor 1 (TNFR1) located in the plasma membrane leads to receptor aggregation and the coordinated recruitment of signaling molecules and the formation of two signaling complexes (*23*). Signaling complex I forms directly at TNFR1 and stimulates pro-survival NF-κB signaling, whereas induction of the extrinsic apoptosis pathway is initiated by complex II formation in the cytoplasm (*24*). While different mechanisms have been shown to induce formation of complex IIa or complex IIb (*23*), several proteins are present in both complexes, including FADD, RIPK1 and pro-caspase 8, which initiates the extrinsic apoptotic cascade. NF-κB activation leads to Cellular FLICE (FADD-like IL-1β-converting enzyme)-inhibitory protein (cFLIP) production, which in turn inhibits caspase-8 cleavage (*25*). We have previously shown that BETi transcriptionally regulate intrinsic apoptotic regulators (*26*), therefore the dependency of these divergent apoptotic pathways downstream of TNF was assessed. HCT-116 and MKN45 cells treated with TNF demonstrated enhanced activation of caspase-8 in the presence of RG6146 (Figure 3A and Supplementary Figure 3A), and downstream activation of caspases-3/7 was suppressed with a specific irreversible inhibitor of caspase-8, Z-IETD-fmk (Figure 3B and Supplementary Figure 3B). Furthermore, siRNA-mediated knockdown of caspase-8 in HCT-116 cells partially rescued the effect of TNF and RG6146 in co-treated cells (Figure 3C and Supplementary Figure 3C and Supplementary Video 3). In support of these findings, ectopic overexpression of cFLIP (an inhibitor of death receptor-mediated apoptosis) was sufficient to protect MC38 cells from the cytotoxic effects of TNF alone and in combination with RG6146 (Figure 3D). Meanwhile, ectopic overexpression of Bcl-2, which primarily regulates the intrinsic apoptotic pathway, was unable to rescue MC38 cells from the cytotoxic effects of TNF, highlighting a clear role for the extrinsic apoptotic pathway mediating the primary cytotoxicity in response to TNF and BETi (Figure 3D). As a functional downstream surrogate of caspase activation, RG6146 and other BETi were capable of enhancing caspase substrate cleavage induced by TNF, as evidence by PARP cleavage in HCT-116, MKN45, and MC38-Ova cells (Figure 3E-F and Supplementary Figure 3D-F). Together, these data demonstrate that BETi sensitize tumor cells to the cytotoxic effects of TNF through enhanced activation of the extrinsic apoptotic cascade.

**Figure 3.**
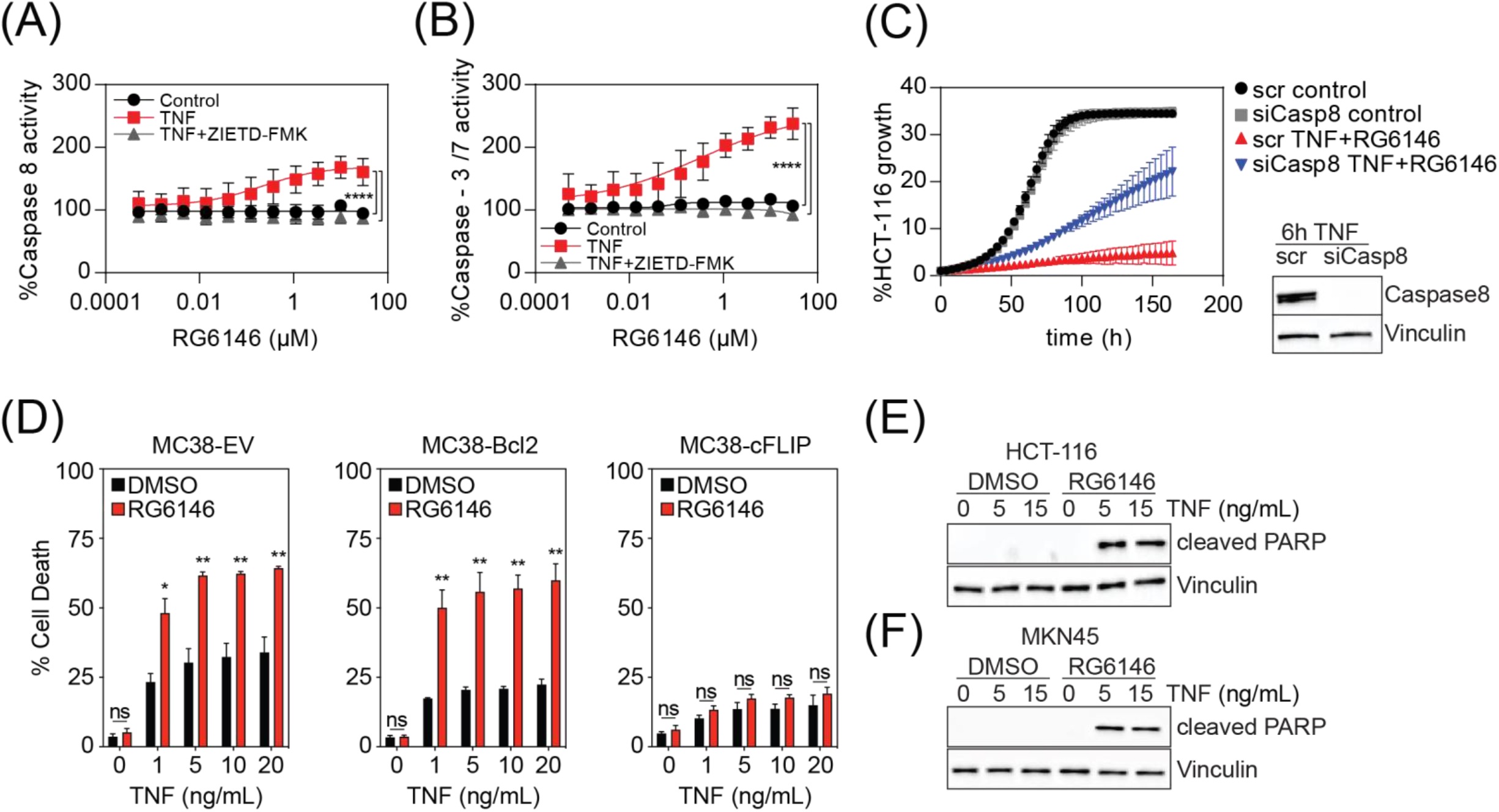
BET inhibition promotes hyperactivation of the extrinsic apoptotic cascade. HCT-116 cells were treated with increasing concentrations of RG6146 in the presence or absence of TNF (15ng/mL) for 8 hours prior to assessment of **(A)** Caspase-8 and **(B)** Caspase-3/7 activity by Caspase-Glo. Caspase-8 inhibitor (1μM ZIETD-FMK) was used as a negative control. Data is normalized to each DMSO control and represents mean of three independent experiments. Statistical significance denotes difference between highest concentration of RG6146 single agent vs combination with recombinant human TNF +/- ZIEDT-FMK (2-way ANOVA, ****p<0.0001). **(C)** HCT-116 cells were reverse transfected with siPOOLs targeting Caspase-8 or a scrambled (scr) control prior to treatment with recombinant human TNF (15ng/mL) and RG6146 (2.5μM) or control. Cell growth was monitored over time by detecting cell confluence using live cell imaging. Knockdown efficacy was detected by western blot of transfected cells after 6h treatment with 15ng/mL TNF. Results were verified in three biological independent experiments. Data of one representative experiment is shown. **(D)** MC38 cells expressing ectopic murine Bcl-2 or cFLIP (CFLAR) cDNA, or an empty vector (EV) control, were exposed to increasing concentrations of recombinant murine TNF in the presence of RG6146 (2.5µM) or DMSO for 24 hours prior to assessment of tumor cell death by flow cytometry (propidium iodide positivity). Data represents mean of three independent experiments. Statistical significance denotes difference between DMSO and RG6146 at each dose of TNF (2-way ANOVA, *p<0.05, **p<0.01). Immunoblot analysis of **(E)** HCT-116 and **(F)** MKN45 cells treated in the presence or absence of RG6146 (2.5μM) and recombinant human TNF (5-15ng/mL) for 24 hours and detection of cleaved PARP and Vinculin.

### BET proteins are required for transcription of pro-survival NF-κB target genes

Concomitant with activation of the extrinsic apoptotic cascade, TNF simultaneously activates pro-survival NF-κB signaling (*27*). As BETi function primarily to disrupt RNA polymerase II-driven transcription, the effect of BETi on pro-survival NF-κB signaling downstream of TNF stimulation was investigated. RNA sequencing (RNA-Seq) in distinct human and mouse tumor lines stimulated with TNF in the presence or absence of RG6146 was performed. As expected, TNF-treated MKN45 and HCT-116 cells demonstrated upregulation of canonical NF-κB target genes (Figure 4A and Supplementary Figure 4A) and co-treatment with RG6146 suppressed the TNF-induced activation of a subset of these genes, while other cluster(s) of genes induced following TNF treatment were less affected by co-treatment with RG6146 (Figure 4B-D and Supplementary Figure 4B-E). The ability of BETi to selectively reprogram the transcriptional response to TNF was also observed in E0771 and MC38 mouse tumor cell lines (Supplementary Figure 4F-G). Finally, we also re-analyzed published expression profiling in non-transformed endothelial cells (*28*) and found prototypical BET inhibitor JQ1 could selectively suppress a subset of TNF stimulated genes that were bound by NF-κB (Supplementary Figure 5A-C). Taken together, these findings suggest a role for BET proteins in the activation of NF-κB signaling following inflammatory stimuli, such as TNF, that is conserved evolutionarily. Among the TNF-induced NF-κB target genes transcriptionally suppressed by RG6146 were important negative regulators of death receptor-induced apoptosis, including *BIRC2* (cIAP1), *BIRC3* (cIAP2), and *TRAF1* (Figure 4E), which also showed suppression at the level of protein expression (Figure 4F and Supplementary Figure 5D). Indeed, ectopic overexpression of both cIAP1 and cIAP2 rescued the combinatorial apoptotic effect of TNF and RG6146 (Figure 4G and Supplementary Figure 5F). Overexpression of cIAP1 alone did not affect the combined effects of TNF and RG6146, while overexpression of cIAP2 had a partial effect (Supplementary Figure 5E-F). These data illustrate that BET inhibition suppresses NF-κB target genes following TNF stimulation to subvert pro-survival signaling, resulting in preferential apoptosis induction and highlights the functional importance of cIAP1 and cIAP2 in regulating death receptor-mediated apoptosis.

**Figure 4.**
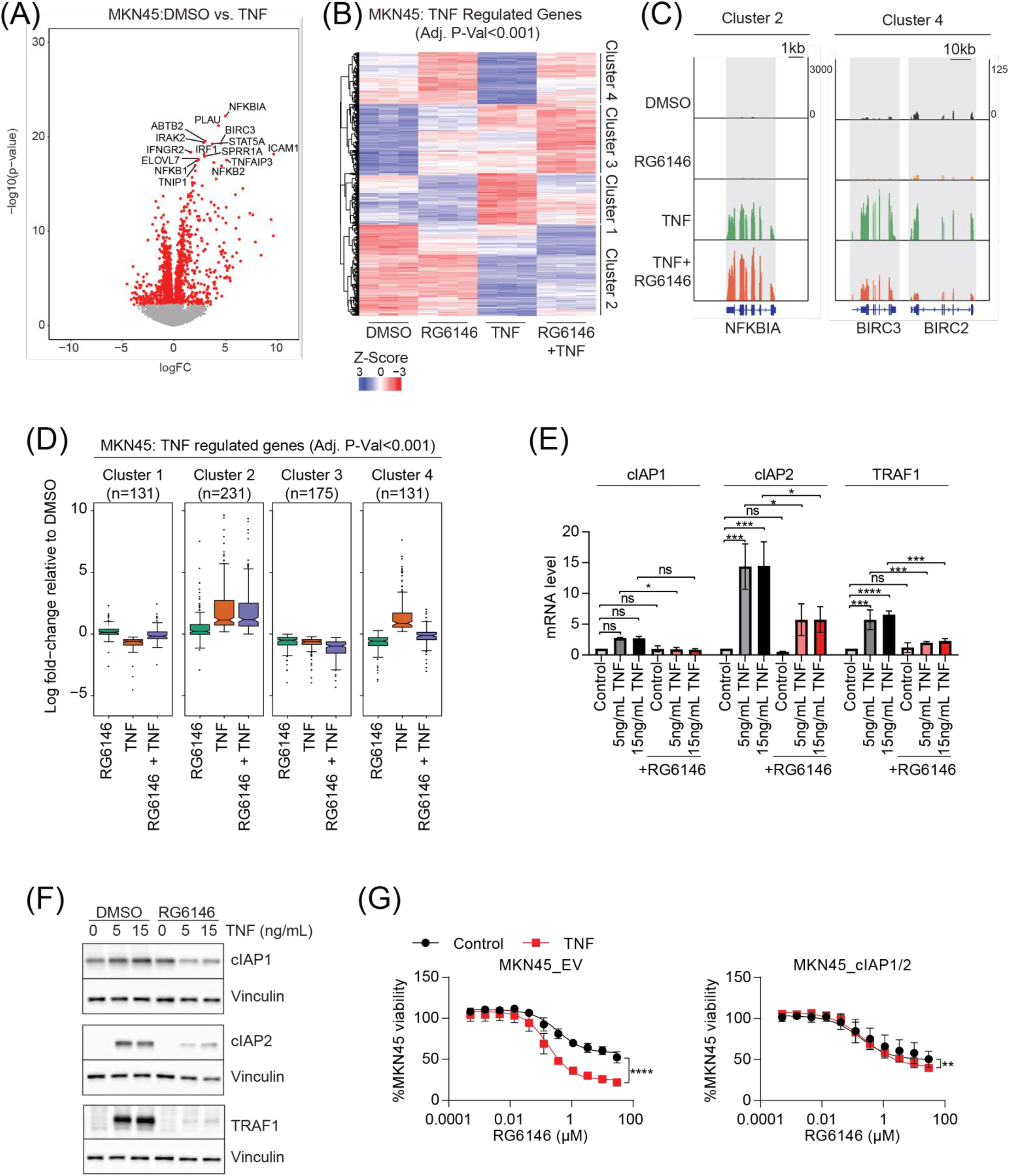
TNF stimulates a BET protein-dependent NF-kB transcriptional response. **(A)** RNA-seq from MKN45 cells treated in triplicate with recombinant human TNF (20 ng/mL) or DMSO vehicle. Differentially expressed genes (red) are defined as adjusted P-value<0.001. **(B)** Heatmap of normalized log counts per million (CPM) of genes differentially expressed in response to TNF (adj. P-value<0.001) separated into 4 groups by k-means clustering. **(C)** IGV genome browser screenshot showing RNA-seq signal at NFKBIA (representative Cluster 2 gene) and BIRC2/3 loci (representative Cluster 4 loci). **(D)** Boxplot of log fold-change values comparing RG6146, TNF, or the combination of RG6146 and TNF, with DMSO for genes within each of the 4 clusters. **(E)** qRT-PCR of MKN45 cells following 2 hours of treatment with recombinant human TNF in the presence or absence of RG6146 (2.5µM) to detect mRNA levels of *BIRC2* (cIAP1), *BIRC3* (cIAP2) and *TRAF1*. Data is normalized to *GAPDH* and represents mean of three independent experiments. Statistical significance denotes difference between TNF and control as well as each condition +/- RG6146 (1-way ANOVA, *p<0.51, ***p<0.001, ****p<0.0001). **(F)** Immunoblot analysis of cIAP1, cIAP2, and TRAF1 protein in MKN45 cells following 24 hours of treatment with recombinant human TNF in the presence or absence of RG6146 (2.5µM). Vinculin was used as a loading control. Data is one representative blot of three independent experiments performed. **(G)** MKN45 cells ectopically overexpressing both cIAP1 and cIAP2 (MKN45_cIAP1/2) or empty vector control (MKN45_EV) were treated with increasing concentrations of RG6146 in the presence or absence of recombinant human TNF (15ng/mL) for 72 hours prior to assessment of viability by CTG2.0. Data is normalized to DMSO and represents mean of three independent experiments. Statistical significance denotes difference between highest concentration of RG6146 single agent and combination with TNF (2-way ANOVA, **p<0.01, ****p<0.0001).

### RG6146 inhibits the recruitment of Brd4 to p65-bound cis-regulatory elements

To understand the mechanistic basis of the transcriptional consequences, epigenomic profiling was performed to explore the ability of RG6146 to modulate the expression of NF-κB target genes. First, chromatin immunoprecipitation and sequencing (ChIP-seq) was performed using MC38 cells to define the genomic binding sites of p65 that were uniquely induced following TNF stimulation (Figure 5A). These analyses also revealed that RG6146 co-treatment was also associated with a small but significant reduction in p65 binding. To understand the differential usage of these p65-bound *cis*-regulatory elements, assay for transposase accessibility (ATAC-seq) was performed to assess differential chromatin accessibility in the presence and absence of TNF. This demonstrated that TNF stimulation resulted in rapid increases in DNA accessibility, suggesting that TNF-induced recruitment of p65 to chromatin is associated with dynamic chromatin remodeling (Figure 5B). Moreover, multiple p65/NF-κB and core promoter (CAAT box)-binding NFY transcription factor (TF) motifs were among the most significantly overrepresented TF motifs in differentially accessible regions of DNA between TNF-stimulated and control cells (Supplementary Figure 6A). Finally, ATAC-seq revealed that neither baseline accessibility nor the increased accessibility in response to TNF stimulation was affected by acute loss of BET protein binding (Figure 5B).

**Figure 5.**
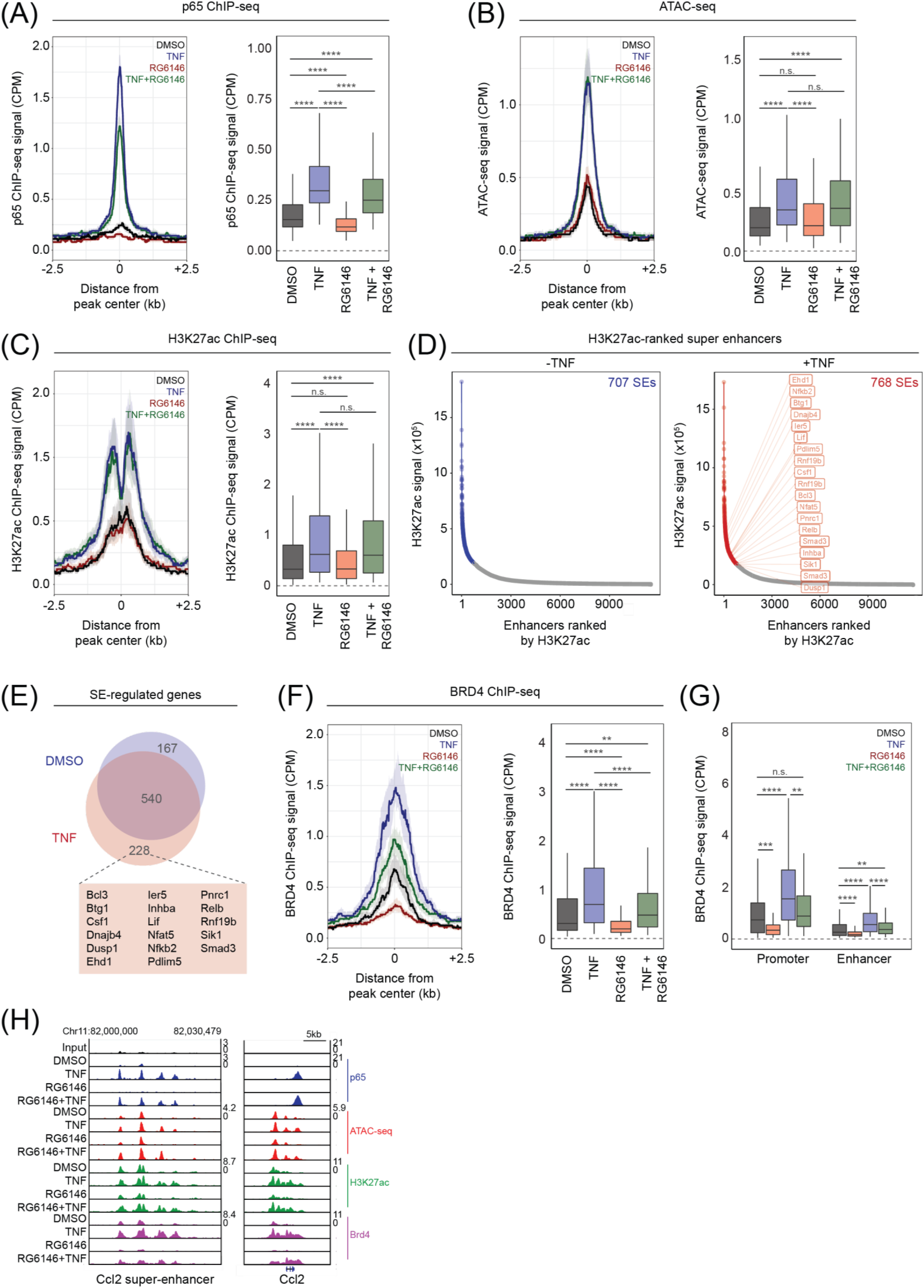
TNF stimulates acute epigenetic remodelling of p65-bound *cis*-regulatory elements. **(A)** Average profile of normalized p65 ChIP-seq signal (counts per million, CPM) at top 350 sites (± 2.5kb) where p65 is recruited following TNF stimulation of MC38 cells treated for 3 hours with RG6146 (2.5µM), TNF (10ng/mL), or the combination. Boxplot represents quantification of p65 ChIP-seq signal across these regions. **(B)** Average profile of normalized ATAC-seq signal (CPM) at p65-bound sites ± 2.5kb (*from A*). Boxplot represents quantification of ATAC-seq signal across these regions. **(C)** Average profile of H3K27ac ChIP-seq signal (adjusted counts per million, CPM) at p65-bound sites ± 2.5kb (*from A*) in MC38 cells. Boxplot represents quantification of H3K27ac ChIP-seq signal across these regions. **(D)** H3K27ac signal at super-enhancers ranked by H3K27ac ChIP-seq signal from MC38 cells treated with DMSO or TNF (10ng/mL). **(E)** Venn diagram overlap of genes regulated by SEs in the presence or absence of TNF, highlighting those genes that overlap with MGSigDB’s ‘TNF via NF-kB’ gene signature. **(F)** Average profile of BRD4 ChIP-seq signal (adjusted counts per million, CPM) at p65-bound sites ± 2.5kb (*from A*) in MC38 cells. Boxplot represents quantification of BRD4 ChIP-seq signal across these regions. **(G)** BRD4 ChIP-seq signal at p65-bound sites classified as either promoter (±5k.b. from annotated TSS) or enhancer (>5k.b. from annotated TSS). **(H)** IGV genome browser screenshot showing ChIP-seq signal at the Ccl2 locus and upstream SE.

In addition, analysis of H3K27 acetylation by ChIP-seq revealed that p65-bound sites were rapidly hyperacetylated in response to TNF stimulation (Figure 5C). Indeed, activation of gene expression by p65 is dependent upon physical co-recruitment of histone acetyltransferases P300/CBP in certain contexts (*29*). These analyses also demonstrated that TNF-induced histone acetylation was unaffected by concomitant BET inhibition, indicating that BET inhibition modulates the transcriptional cascade downstream of histone acetylation (Figure 5C), which fits a model whereby BET proteins are recruited to acetylated nucleosomes/TFs. In parallel, super enhancers demarcated by H3K27ac were assessed and revealed that TNF stimulation leads to dynamic changes in super enhancer landscape (Figure 5D), where many canonical NF-κB regulated genes gain SE regulatory elements following TNF stimulation (Figure 5E). Finally, the ability for RG6146 to modulate genome-wide occupancy of Brd4 was assessed. Under steady state conditions, acute RG6146 exposures led to a profound loss of Brd4 occupancy genome-wide (Supplementary Figure 6B), including displacement of Brd4 from H3K27ac-ranked super enhancers and typical enhancers (Supplementary Figure 6C-F). In the context of TNF stimulation, RG6146 was capable of partially abrogating the recruitment of Brd4 to p65-bound *cis*-regulatory elements (Figure 5F), consistent with observations from RNA-seq analysis that selective suppression of transcription is observed rather than complete suppression. Moreover, significant loss of Brd4 binding was observed at both promoter (peak ≤5kb from annotated TSS) and *cis*-regulatory enhancer elements that drive the transcription of p65 target genes (Figure 5G), such as the SE-driven chemokine *Ccl2* (Figure 5H). Overall, these findings suggest that RG6146 functions to abrogate the transcriptional response to TNF by limiting the *de novo* recruitment of BRD4 to *cis*-regulatory elements that control inflammatory gene expression.

### BET inhibition promotes TNF-dependent bystander killing and increases the anti-tumor activity of T-cell bi-specific antibodies

Active anti-tumor immune responses are associated with secretion of TNF and other pro-inflammatory cytokines into the TME (*21, 30*). Moreover, it is known that IO agents are capable of further increasing the secretion of pro-inflammatory cytokines (*31, 32*). Therefore, the ability of BETi to potentiate the cytotoxic activity of CTL-derived TNF that is induced by IO agents was investigated. T-cell bispecific (TCB) antibodies are IO agents that simultaneously engage with target antigens on T-cells and cancer cells leading to potent antigen-specific T-cell killing (*33, 34*). The 2:1 CEA-TCB antibody is engineered to engage tumor cells expressing high levels of carcinoembryonic antigen (CEA) on their surface, via two CEA-specific Fab regions, with CD3 on the surface of T-cells via one CD3-specific Fab region (*34*). Co-culture of MKN45 cells (which express endogenous CEA) and human peripheral blood mononuclear cells (PBMCs) in the presence of picomolar concentrations of CEA-TCB led to the release of granzyme-B, IFN-γ, and TNF into the culture supernatant and potent anti-tumor activity (Supplementary Figure 7A-B). The role of secreted factors in mediating anti-tumor responses by TCB antibodies was investigated using an assay in which supernatant (SN) derived from a PBMC:MKN45 co-culture treated with CEA-TCB was transferred to fresh HCT-116 or MKN45 tumor target cells (Figure 6A). Exposure to co-culture SN led to potent induction of tumor cell death in both tumor lines, which could be further enhanced by addition of RG6146 (Figure 6B-C and Supplementary Figure 7C-D). Moreover, RG6146 promoted the cleavage of PARP in MKN45 cells following transfer of SN (Figure 6D), indicating the presence of soluble factors capable of inducing tumor cell death. Addition of a TNF-neutralizing antibody rescued the cell death induced by TNF and RG6146 co-treatment (Figure 6E), identifying TNF as the key soluble factor in TCB SN. Taken together, these data indicate that novel 2:1 TCB antibodies promote the secretion of soluble TNF from CD8^+^ T-cells and that BET inhibition can be utilized to increase the sensitivity of tumor cells to CTL-derived TNF in the context of TCB antibody treatment.

**Figure 6.**
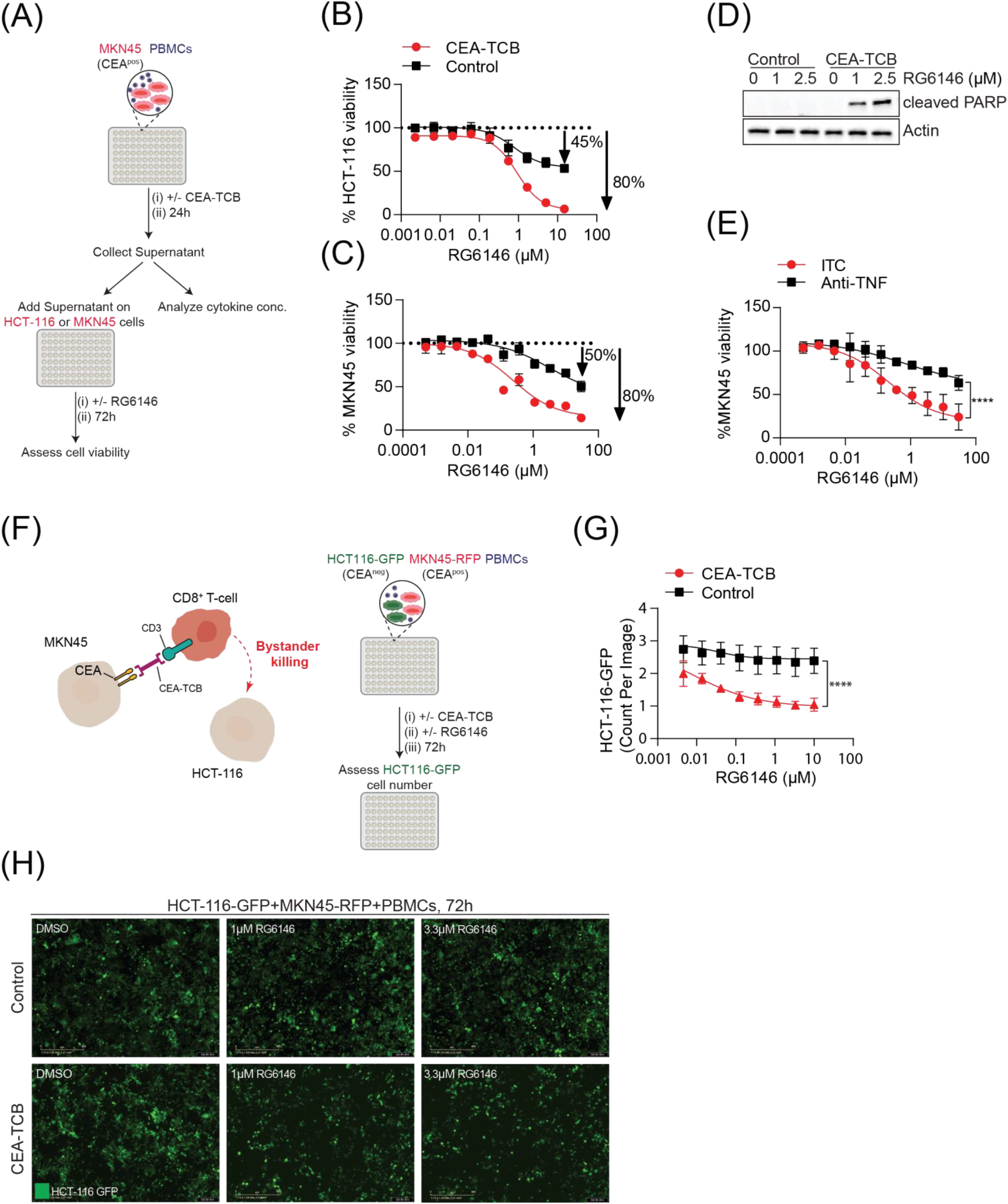
BET inhibition promotes the activity of T-cell bi-specific (TCB) antibodies. **(A)** Schematic of TCB assay where PBMCs co-cultured with MKN45 cells (expressing CEA) are treated for 24 hours with 20nM CEA-TCB or vehicle control. Supernatant was collected and cytokine concentration analyzed by flow cytometry, or added to naïve HCT-116 or MKN45 cells and treated with a dose response of RG6146. Cell viability was assessed after 72h treatment. **(B-C)** Viability of HCT-116 (B) and MKN45 (C) following treatment with supernatant from a co-culture assay treated with 20nM CEA-TCB (CEA-TCB) or control (Control). Data of one representative experiment is shown and was normalized to DMSO control. Results were verified in three independent experiments. **(D)** Immunoblot for cleaved PARP and Vinculin from MKN45 cells following treatment with supernatant from a co-culture treated with 20nM CEA-TCB (CEA-TCB) or control (Control) and 1 or 2.5μM RG6146 for 24 hours. One representative blot of three independent experiments is shown. **(E)** Viability of MKN45 cells following treatment with supernatant from a supernatant co-culture assay treated with 20nM CEA-TCB and 10μg/mL TNF alpha Monoclonal Antibody (Anti-TNF) or isotype control (ITC). Viability was assessed following 72h treatment with increasing doses of RG6146. Data was normalized to DMSO and represents mean of three independent experiments. Statistical significance denotes difference between highest concentration of RG6146 with Anti-TNF or ITC (2-way ANOVA, ****p<0.0001). **(F)** Schematic of bystander killing assay in which PBMCs were co-cultured with MKN45-RFP cells (CEA^+^) and HCT-116-GFP cells (CEA^-^) and then treated with 40nM CEA-TCB alone or in combination with increasing doses of RG6146 for 72 hours. Growth of HCT-116-GFP cells was monitored over time and confluence analyzed after 72 hours. **(G)** Relative cell count of HCT-116-GFP cells normalized to time point of treatment (T0). Data represents mean of three independent experiments. Statistical significance denotes difference between highest concentration of RG6146 in combination with CEA-TCB or control (2-way ANOVA, ****p<0.0001). **(H)** Representative images from by-stander killing assays at 72 hour time point.

Following the recognition of cognate tumor antigen in the context of MHC class I, the delivery of cytotoxic granules is spatially restricted to the immunological synapse. In contrast, pro-inflammatory cytokines can diffuse within the TME to induce paracrine inflammatory responses. As such, TNF could therefore promote death of tumor cells that do not directly interact with CTLs and/or antigen-loss variants through so called ‘bystander killing’. A co-culture assay mimicking bystander killing (Figure 6F) was utilize to determine if RG6146 promoted bystander killing of HCT-116 cells, which do not express endogenous CEA (Supplementary Figure 7E). The growth of HCT-116 cells (expressing GFP) co-cultured with MKN45 cells (expressing RFP) and PBMCs was monitored over time (Figure 6F). In these co-culture assays, addition of CEA-TCB in combination with RG6146 significantly decreased HCT-116 cell number compared to single agent RG6146 (Figure 6G-H). Finally, the ability for BETi to augment antigen-independent bystander killing of tumors was independently validated in an analogous mouse co-culture system using a mixture of antigen (Ova) positive and negative MC38 tumor cells. In this assay, antigen negative MC38 cells were chromium (^51^Cr)-labeled and mixed 1:1 with non-chromium-labeled MC38-Ova cells, prior to co-culture with Prf^-/-^ OTI T-cells in the presence or absence of RG6146, where lysis of antigen negative cells is detected by the presence of ^51^Cr in the co-culture SN. Consistent with data from human cell lines, RG6146 was capable of promoting killing of antigen positive tumor cells as well as bystander killing of tumor antigen negative cells (Supplementary Figure 7F). Overall, these experiments demonstrated that BETi are capable of lowering the cytotoxic threshold to TNF in a variety of tumor types, including bystander killing of antigen-loss variants that are not directly recognized by cytotoxic lymphocytes.

### BET inhibition and checkpoint immunotherapy promote TNF-dependent anti-tumor immunity in vivo

The therapeutic effects of combining BETi with CEA-TCB *in vivo* was assessed using the first-generation BETi, JQ1, due to its advanced pre-clinical development and high potency against murine tumors compared to RG6146. MC38 cells were engineered to express CEACAM5 and orthotopically transplanted into syngeneic recipient mice that were subsequently treated with CEA-TCB, JQ1 or the combination of both agents (Figure 7A). These treatment regimens were well tolerated *in vivo* (Supplementary Figure 8A-B) and single agent JQ1 or CEA-TCB treatment decreased tumor growth by 50% and 60%, respectively, (Figure 7B and Supplementary Figure 8C). Importantly, the combination of JQ1 and CEA-TCB induced more profound tumor regression than single agent treatments (Figure 7B and Supplementary Figure 8C). In addition, neutralization of TNF *in vivo* by antibody depletion abrogated the therapeutic effect of BETi in the combination treatment (Figure 7C and Supplementary Figure 8D). To determine if BETi could similarly augment the activity of additional IO agents against solid tumors, MC38 tumors were transplanted into syngeneic recipient mice and treated with anti-PD1 checkpoint immunotherapy, JQ1 or a combination of both agents (Figure 7D). Single agent treatment modestly decreased MC38 tumor growth, while the combination of both agents further suppressed tumor growth and enhanced survival (Figure 7E-F). These data further demonstrate that systemic administration of BETi can potentiate the activity of multiple IO agents *in vivo*, including TCB and immune checkpoint antibodies.

**Figure 7.**
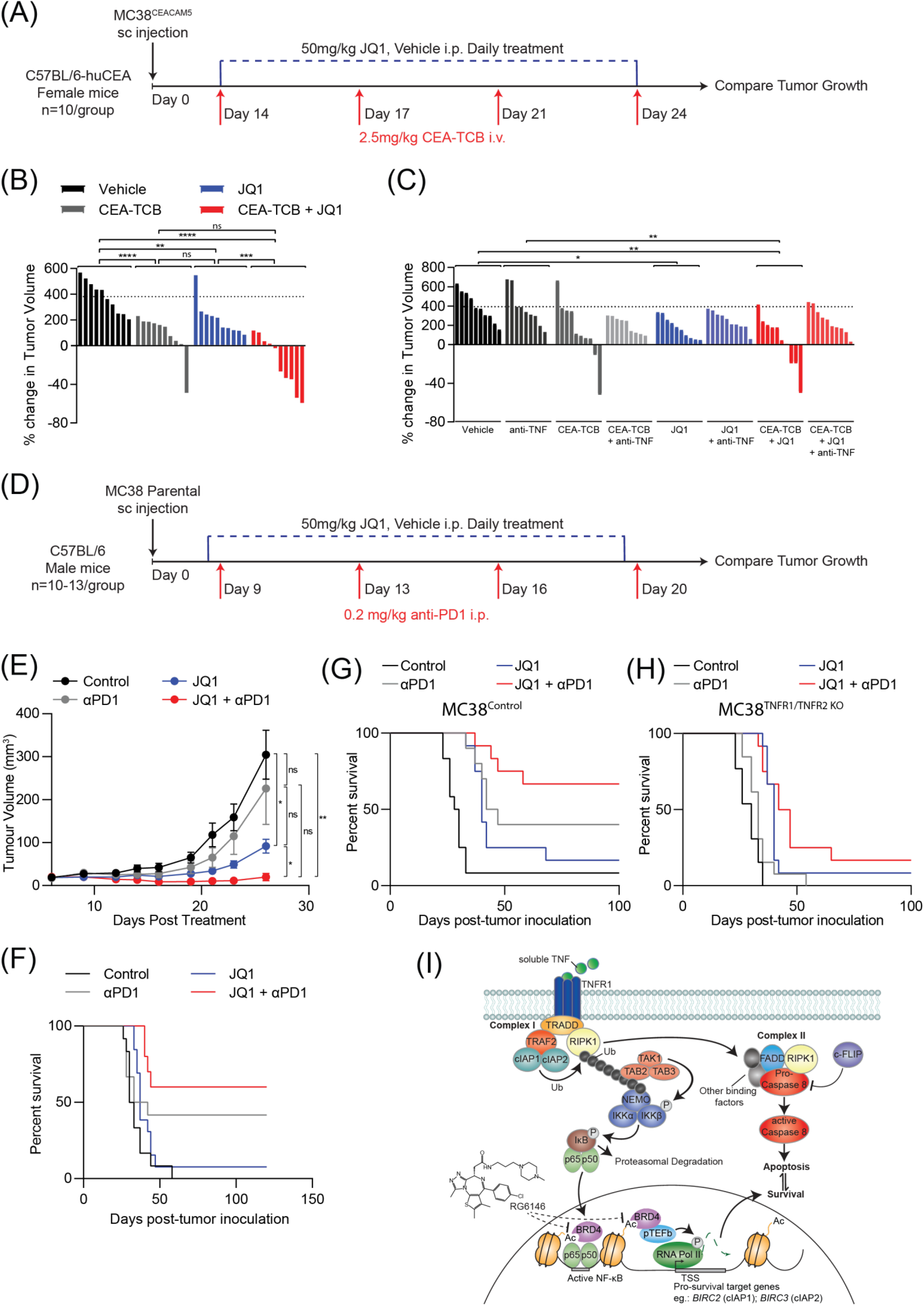
BET inhibition increase the efficacy of immune-oncology agents *in vivo*. **(A)** Schematic of *in vivo* assay. Cohorts of transgenic C57BL/6 mice expressing human CEA (n=10 per treatment group) bearing MC38 tumors engineered to express CEACAM5 were treated with 2.5mg/kg CEA-TCB antibody twice weekly, 50mg/ kg JQ1 daily, or the combination of CEA-TCB and JQ1. **(B)** Bar chart of experiment described in *A* represents change in tumor volume (%) on day 13 compared to day 0 (relative to treatment commencement). Statistical significance denotes difference between treatment groups (2-way ANOVA, **p<0.01, ***p<0.001, ****p<0.0001). **(C)** C57BL/6-huCEA mice bearing MC38-CEACAM5 tumors were treated with JQ1 and CEA-TCB alone and in combination (*as described in A*) and in the presence of a TNF neutralizing antibody twice-weekly. Statistical significance denotes difference treatment groups (2-way ANOVA, *<0.05, **p<0.01; if not indicated not significant). **(D)** Schematic of *in vivo* assay. Wild-type C57BL/6 mice bearing MC38 tumors (n=10-13 mice per treatment group) were treated daily with 50mg/kg JQ1 or DMSO control alone (i.p.) and in combination with 0.2mg/kg anti-PD1 (i.p.) twice weekly. **(E)** Results of experiment described in *D*. Average tumor growth curves show average tumor growth in each treatment condition. Data represents mean of two independent experiments (+/- SEM). Statistical significance denotes difference between treatment groups of the last tumor measurement (2-way ANOVA, *p<0.05, **p<0.01). **(F)** Kaplan-meier survival curve showing the overall survival of mice from the experiment described in *(D)*. **(G-H)** Results of experiment described in *D* except MC38 cells depleted of (G) vehicle control (MC38 control) or (H) TNFR1 and TNFR2 (MC38 TNFR1/2 KO) were injected into mice. Kaplan-meier survival curve showing the overall survival of mice. Data represents mean of two independent experiments. **(I)** Mechanism of BET protein mediated NF-κB signalling downstream of TNF stimulation: Ligation of TNF to the TNF receptor (TNFR1) induces formation of complex I, comprising TRADD, RIPK1, TRAF2 and cIAP1 and cIAP2. Complex I components, cIAP1 and cIAP2 ubiquitinate RIPK1 which leads to the recruitment of additional binding factors that ultimately induce degradation of IκB. Loss of IκB promotes nuclear translocation of active NF-κB transcription factor (p65/p50), which stimulates the expression of NF-κB target genes in BET protein-dependent manner. BET inhibition abrogates the transcription of pro-survival genes, including *BIRC2* and *BIRC3*, limiting pro-survival NF-κB signalling. As a consequence, BET inhibition leads to unrestrained activation of complex II, which includes FADD, RIPK1, and pro-caspase-8, and engagement of the extrinsic apoptotic cascade and increased cell death.

Finally, we investigated the dependence on tumor cell intrinsic TNF signaling for the observed therapeutic responses to BETi and anti-PD1 checkpoint immunotherapy by generating MC38 cells deficient in cognate TNF receptors, TNFR1 and TNFR2. For subsequent use in syngeneic recipients, MC38 cells were engineered by electroporation of Cas9-sgRNA ribonucleoprotein complexes, thereby avoiding the immunogenicity of constitutive Cas9 expression (Supplementary Figure 9A). Individual depletion of TNFR1 (26 days median survival) or TNFR2 (31.5 days median survival) had minimal effects on the growth of tumors in vehicle-treated mice compared to mice bearing control tumors (29 days median survival; Figure 7G and Supplementary Figure 9B-D,F-G). In the context of anti-PD1 checkpoint immunotherapy, we found reduced median survival of mice bearing TNFR1 (33 days median survival) or TNFR2 (40 days median survival) deficient tumors compared to mice bearing control tumors (44.5 days median survival). In addition, the frequency of complete remissions with anti-PD1 were reduced from 40% (in mice bearing control tumors) to 0% and 16.6% for TNFR1 and TNFR2 deficient tumors, respectively, demonstrating a clear role for TNFR1/TNFR2 engagement for responses to immune checkpoint blockade. While the median survival in response to single agent JQ1 therapy was largely unaffected by individual TNFR1 (36 days median survival) or TNFR2 (40 days median survival) depletion (compared to 40 days median survival for mice bearing control tumors), we noted the frequency of complete remissions were reduced from 16.6% (in mice bearing control tumors) to 0% for both TNFR1 and TNFR2 deficient tumors, respectively (Figure 7G and Supplementary Figure 9B-D,F-G). Finally, we found that dual depletion of both TNFR1 and TNFR2 had no effect on tumor growth in vehicle treated mice (30 days median survival) and a modest reduction in survival from anti-PD1-treatment (33 days median survival). More strikingly, we noted dual TNFR1/TNFR2 depletion significantly reduced the efficacy of BETi and anti-PD1 combination therapy (44.5 days median survival) for which completed remissions were limited to 16.7% compared to 66.7% in the control group (Figure 7G-H and Supplementary Figure 9B-E). Taken together, these data demonstrate the key physiological role for TME-derived TNF and tumor cell intrinsic TNF signaling in mediating therapeutic response to BETi and IO agents *in vivo*. Moreover, these findings further highlight the immune modulatory capacity of BETi and validate an additional immune-dependent mechanism by which BETi evoke anti-cancer responses *in vivo*.

## DISCUSSION

Signal transduction pathways activated by pro-inflammatory cytokines are well characterized, however, it is also clear that active epigenetic remodeling is a critical prerequisite for coordinated inflammatory gene expression. This study highlights a concerted epigenetic dependency on BET proteins for NF-κB activation in response to TNF stimulation. Dynamic redistribution of BRD4 was required for the rapid activation of NF-κB target genes and protection of tumors from CD8^+^ T-cell killing by TNF. By subverting the recruitment of BRD4 to p65-bound *cis*-regulatory elements, BET inhibition broadly potentiated the cytotoxic activity of TNF across cancer cell lines of diverse tissue origin. Key negative regulators of the extrinsic apoptotic cascade including cIAP1 and cIAP2 were directly suppressed by BETi, and overexpression of these anti-apoptotic factors could partially rescue the phenotype observed with RG6146 and TNF. These genetic experiments are consistent with previously published findings that genetic manipulation of individual genes are largely unable to recapitulate the complex signaling associated with TNF biology (*35*). Thus, rather than attributing the cooperative effects of BETi on TNF-mediated cell killing to the modulation of a single gene, these effects likely result from transcriptional modulation of multiple TNF-regulated genes simultaneously. The ability of BETi to sensitize solid tumor cells to the TNF induced apoptosis was not restricted to antigen expressing tumor cells directly engaged by T-cells, but also resulted from enhanced bystander killing of tumor cells independently of direct antigen recognition. By simultaneously increasing TNF-induced tumor cell death and suppressing PD-L1 expression (*16*), BETi exhibit multiple favorable immunomodulatory properties as potential IO and CART adjuvants.

The physiological role of BET proteins has been extensively studied in a variety of non-malignant pathologies. The first reported BETi was shown to selectively reprogram transcriptional responses to cytokines in the context of acute inflammation (*28, 36*), where transcriptional suppression of TNF-inducible pro-inflammatory cytokines was observed in LPS-stimulated bone marrow-derived macrophages. Importantly, it has been reported that BET proteins bind directly to acetylated p65 leading to stabilization of active NF-κB (*37*). These studies, and others, have led to the suggestion that BETi may be developed therapeutically as anti-inflammatory agents for chronic inflammatory diseases. Given BET proteins are ubiquitously expressed, our findings raise the possibility that if BETi are deployed during acute inflammation, healthy tissue proximal to the site of inflammation may inadvertently be sensitized to the cytotoxic effects of TNF, highlighting the potential for exacerbation of autoimmunity by BETi. Consistent with RNA-seq assays performed in tumor specimens, reanalysis of expression profiling in primary human epithelial cells treated with TNF and JQ1 revealed selective modulation of transcription response by concurrent BET inhibition, although no assessment of cell death was reported in that study. Thus, careful examination for pathophysiological signs of autoimmunity are warranted where BETi are currently being utilized clinically. Importantly pre-clinical studies using syngeneic solid tumor models showed that combination regimens incorporating BETi and IO agents were well tolerated and lead to enhanced anti-tumor responses.

While profound cytotoxicity in response to BETi and exogenous or CTL-derived TNF was demonstrated in a variety of cell lines, this mechanism of immunomodulation was not observed in cell lines and patient specimens that were inherently refractory to the cytotoxic effects of TNF. Resistance to the cytotoxic effects of TNF are incompletely understood, but have been previously linked to altered expression or mutation in genes encoding requisite TNF pathway members, disruption of wild-type p53 function, cleavage of second messenger molecules, and constitutive activation of the NF-κB pathway (*38-43*). In order to effectively exploit this immunomodulatory effect of BETi clinically, the identification of predictive biomarkers would be beneficial to define patient populations responsive to TNF. Thus, ongoing functional genomics studies aim to identify genetic, epigenetic, or transcriptomic features that distinguish those highly TNF responsive versus non-responsive tumors.

Overall, these data support a model where pro-survival NF-κB signaling is BET protein-dependent and protects tumor cells from CD8^+^ T-cell-derived TNF during anti-tumor immune responses. Importantly, this mechanistic dependency is therapeutically exploitable where BET inhibitors selectively perturb the transcription of pro-survival NF-κB target genes and potently sensitize tumor cells to the cytotoxic effects of TNF (Figure 7I). Pre-clinical experiments with TCB and anti-PD1 antibodies implicated this novel mechanism in the augmented therapeutic IO responses *in vivo*. These results provide new insights into the targeting of BET proteins and demonstrate that disruption of pro-inflammatory cytokine responses by epigenetic reprogramming is a novel mechanism of therapeutic immunomodulation in cancer.

## Author Contributions

S.J.H. and L.C.W. developed the concepts, performed and analyzed experiments, and wrote and edited the manuscript. Additional experiments were performed and analyzed by D.N.M., T.F. and D.G. (*in vivo* assays), P.T., L.A.C, C.J.K and H.J. (in *vitro assays*), K.M.R. (live cell imaging), J.M. (tumoroid cultures). J.O. provided guidance on live cell imaging and tumoroid cultures. M.B and T.F. provided guidance on TCB assays. S.J.V, A.P., D.M. and L.J. assisted with data analysis. J.S. assisted with study supervision. D.R., A.R-B. and R.W.J. developed the concepts and overall strategy, wrote and edited the manuscript.

## Acknowledgements

We thank Dr. Vignesh Narasimhan and Prof. Robert Ramsay (Peter MacCallum Cancer Center, Melbourne, Australia) for assistance with patient-derived colorectal tumoroid cultures. We thank Prof. John Silke (Walter and Eliza Hall Institute, Melbourne, Australia) and members of the Johnstone Laboratory for helpful discussions. The laboratory of R.W.J. was supported by the Cancer Council of Victoria, National Health and Medical Research Council of Australia (NHMRC) and The Kids’ Cancer Project (R.J.W.). S.J.H. was supported by a Postdoctoral Fellowship from the Cancer Council of Victoria. The Peter MacCallum Foundation and Australian Cancer Research Foundation provide generous support for equipment and core facilities.

## MATERIAL AND METHODS

### Cell lines

HCT-116 and MKN45 cell lines and derivatives thereof were cultured at 37ºC and 5% CO_2_ in RPMI-1640 ATCC formulation Medium (#A10491) supplemented with 10% FBS (#10270106) and RPMI supplemented with GlutaMax (#61870) and 20% heat inactivated FBS (hiFBS), respectively. All human cell lines were provided by the Roche Non-Clinical Biorepository from Roche Basel or the Roche-Innovation Center Zurich (RICZ). Murine MC38, AT3, E0771 and derivatives thereof, were cultured at 37°C and 10% CO_2_ in Dulbecco’s modified Eagle’s medium (DMEM) supplemented with 10% FBS and penicillin/streptomycin (Gibco). Peripheral Blood Mononuclear Cells (PBMCs) were isolated from Buffy Coats provided by the Blutspendezentrum SRK beider Basel using density gradient separation (Ficoll-Paque Premium, Sigma #GE17-5442-02). CMV-specific T-cells were kindly provided by RICZ as previously described (*44*) and were expanded every four weeks. Freshly isolated PBMCs were kept in a cell culture flask for 1 hour to separate adherent monocytes. Lectin from Phaseolus vulgaris (Sigma #L2796) was added for 1 hour to the cell solution (Feeder cells). LCL cells were loaded with 10nM CMV pp65 peptide NLVPMVATV (NLV) (Thinkpeptides) and incubated for 1 hour. LCL-NLV and Feeder cells were irradiated (LCL-NLV cells: 5000 rad, Feeder cells: 2500 rad). 10,000 CMV-specific T-cells were mixed with LCL-NLV and Feeder cells (ratio 1:5:125) in a 96-well plate in RPMI + GlutaMax + 10% hiFBS + 400U/ml IL2 (Proleukin Novartis) and expanded when T cells started to proliferate

### Compound preparation

RG6146 (Roche), (+)-JQ1 (#S7110), ABBV-744 (#S8723), Mivebresib (#S8400), OTX015 (#S7360), Apabetalone (#S7295), I-BRD9 (#S7835), GSK J4 HCl (#S7070), C646 (#S7152), SGC707 (#S7832), EPZ-6438 (#S7128), SGC 0946 (#S7079), EPZ015666 (#S7748), Birinapant (#S7015), LCL161 (#S7009) all Selleck Chemicals, I-CBP112 (BPS Bioscience #27336), SGC-CBP30 (#T6668), Entinostat (#T6233), Panobinostat (#T2383), Vorinostat (#T1583) all TargetMol, Decitabine (TOCRIS #2624) and I-BET151 (ChemScene #CS-0930) were resuspended to 10-20mM stock concentration in DMSO. iBET-762, Y803, and dBET1 were provided by James E. Bradner (Harvard Medical School). All additional compounds were obtained from Sigma-Aldrich or Selleck Chemicals.

### HCT-116-NLV co-culture-Screen

HCT-116 cells were incubated for 1 hour with 10nM NLV or GLCTLVAML (EBV) peptide (Thinkpeptides), respectively, with rotation and plated for 2 hours at a density of 0.1×10^5^ cells/well in a 96-well plate. CMV-specific T-cells were added (ratio 1:1) and the co-culture was incubated for 30 minutes. Small molecules were added to a final concentration of 2.5-5μM, or DMSO control and cells were incubated for 48 hours before T-cells were removed. Viability was assessed using 100μL CellTiterGlo2.0 (CTG2.0 Promega #G9242). In addition, 20μg/mL TNF-alpha Monoclonal Antibody (28401) (ThermoFisher #MA5-23720) or Mouse IgG1 Isotype Control (MOPC-21) (ThermoFisher #MA1-10407) were added to the co-culture where indicated.

### Activation of OVA-specific OTI T-cells

Whole spleens from 6-10 week old perforin-wild-type (C57Bl/6.OTI) or perforin-deficient (C57Bl/6.OTI.Prf^-/-^) mice were manually dissociated through a 70µM cell strainer and OTI T-cells were activated and expanded with 20ng/mL of SIINFEKL peptide (Sigma-Aldrich) and 100 IU/mL recombinant human IL-2 (Biolegend) in RPMI media supplemented with 10% FBS, GlutaMax (2mM), penicillin/streptomycin, non-essential amino acids, sodium pyruvate (1mM), HEPES (10mM) and 2-mercaptoethanol (50µM). OTI cells were incubated for three days at 37ºC with 5% CO_2_ before being passaged into fresh media (IL-2 only, no SIINFEKL), and cultured for1-4 days prior to use in co-culture killing assays.

### MC38-Ova and OTI co-culture screen

MC38 cells stably expressing MSCV-OVA-GFP (MC38-OVA) were plated at 1.5 × 10^5^ cells per well of a 48-well plate and allowed to adhere for 6-8 hours. Activated OTI cells were washed with PBS and re-suspended in supplemented DMEM prior to serial dilution and addition to MC38 target cells. Cultures were subsequently incubated at 37ºC with 10% CO_2_ for 18-20 hours prior to assessment of induction of tumor cell death by flow cytometry. Co-culture supernatant containing OTI T-cells and dead cells were collected from each well and surviving, adherent MC38 cells were harvested using trypsin. Total well contents were washed in flow cytometry buffer (PBS + 2% FCS + 5mM EDTA) and OTI cells stained with anti-mouse CD5 APC-conjugated antibody (clone 53-7.3; eBioscience). Samples were washed twice with flow cytometry buffer and propidium iodide (PI; 2µg/mL) was added immediately prior to analysis. Analysis was performed on an LSR II flow cytometer (BD Biosciences) and data were analyzed with FlowJo (Tree Star). CD5+ OTI cells were gated out of analysis and dead MC38-OVA cells were recognized as GFP-PI+. Anti-TNF neutralizing antibody (#506325) was obtained from Biolegend.

### Co-culture time lapse microscopy

MC38-OVA cells were seeded into each well of an eight-well chamber slide (Ibidi, Munich, Germany) and incubated overnight at 37°C and 10% CO_2_. Activated OTI T-cells labeled with CellTrace™ Violet (ThermoFisher, #C34557) and re-suspended in supplemented DMEM prior to being added to adherent targets (at a 2:1 tumor to T-cell ratio), in medium containing active caspase-3/7 detection reagent (ThermoFisher). Chamber slides were mounted on a heated stage within a temperature-controlled chamber maintained at 37°C, and constant CO_2_ concentrations (5%) were infused using a gas incubation system with active gas mixer (“The Brick”; Ibidi). Optical sections were acquired through sequential scans or brightfield/differential interference contrast on a TCS SP5 confocal microscope (Leica Microsystems, Deerfield, IL) using a ∼40 Å (numerical aperture, 0.85) air objective and Leica LAS AF software. Image analysis and enumeration of active caspase-3/7 positivity was performed using MetaMorph Imaging Series 7 software (Universal Imaging, Downingtown, PA).

### Cell line screening for sensitization to TNF killing

Cell line screening was outsourced to Oncolead (Karlsfeld, Germany). Briefly, cell lines were obtained from ATCC, NCI, CLS and DSMZ and cultured in recommended medium. Cells were treated with a dose response of RG6146 (10-0.0032μM) or control and 15 ng/mL TNF for 120 hours. Total protein concentration was used to assess the cell number. Adherent and suspension cells were fixed with 10% or 50% tricholoracetic acid, respectively, by incubation for 1 hour at 4 ºC, prior to washing with deionized water and dried. Cell staining was then performed for 30 minutes with 0.04% (wt/v) Sulforhodamine B (SRB) at room temperature. Cells were washed 6x with 1% (v/v) acetic acid and plates were dried before SRB was solubilized in 10mM Tris base prior to measurement of fluorescence with a Seelux-LED96 plate reader (492, 520 and 560nm). Background density (medium only) was subtracted from each experimental well. The TNF synergy was determined by subtracting maximal growth inhibition of RG6146 alone from maximal growth inhibition of RG6146+TNF.

### Murine solid tumor cell death analysis

For cell death analyses of adherent murine solid tumors cells (MC38, AT3, and E0771) and genetically modified variants thereof, were seeded (1.5×10^5^ cells/well) into 48-well plates for ≥8 hours prior to addition of recombinant TNF and BET inhibitors or DMSO/PBS control. Following addition of BET inhibitors (RG6146, JQ1, IBET151, IBET762, dBET 1μM; RVX-208 10μM) in the presence or absence of TNF (5ng/mL), MC38-Ova cells were incubated for 18 hours. Cells were harvested by centrifugation, washed once in ice-cold flow cytometry buffer (2% FBS in PBS), prior to being resuspended in flow cytometry buffer containing PI and assessed for PI positivity. Data were collected on a FACSCanto II flow cytometer (BD Biosciences) and analyzed using FlowJo Software (Version 10.2, Tree Star).

### Synergy Score

Cell death data (where serial dilutions of RG6146 and TNF were used) was imported into Rstudio (Version 3.3.1) and raw values were first visualized using pheatmap package (1.0.12). Data were analyzed using the Synergyfinder package (Version 3.8). Synergy analysis was performed by first calculating a four-parameter log-logistic model to generate the dose-response curves for each single agent. Drug synergy scoring was then calculated using the ZIP algorithm (*45*).

### CRC Organoid microscopy

Matrigel embedded tumors were generated as previously described (*46*). A total volume of 350μL supplemented OB media was applied to the Matrigel-embedded tumoroids. Propidium iodide (PI; Sigma-Aldrich) was added to a final concentration of 2µg/mL. Wells were set up with the following conditions: tumoroids + DMSO vehicle control, tumoroids + recombinant TNF (10ng/mL), tumoroids + RG6146 (2.5µM), and tumoroids + TNF + RG6146. Tumoroids were incubated at 37°C, 5% CO_2_ for 24 hours, and then end-point images were acquired. U-bottom 96-well plates were mounted on a heated stage in a temperature controlled chamber, maintained at 37°C, 5% CO2 (“The Brick”; iBidi). Using the cellSens software (Olympus), imaging of the plate was obtained on an UPLSAPO 4X (NA 0.16) air objective using a Hammamatsu ORCA-Flash 4.0 camera. Z-stack images were acquired by taking 25 sequential images (Z-spacing 50 µm) of PI (emission 656 nm, exposure 368.644 ms) through the matrigel, and Z stack images converted to one image in EFI format. Image quantitation was conducted using Meta-Morph Imaging Series 7 Software. An identical region of interest was identified within the matrigel and integrated morphometry analysis was used to filter out individual T-cells based on size. A threshold was applied to delineate PI positive pixels, and integrated intensity was measured. Identical conditions were applied to compared images.

### Caspase activity assay

Cells were plated at a density of 0.04×10^6^ cells/well in a 96-well plate. After 24 hours cells were treated with a dose response of RG6146, 15ng/mL recombinant human TNF (BioLegend #570104) and 1μM Caspase-8 Inhibitor Z-IETD-FMK (R&D Systems #FMK007) or the corresponding controls (0.15% DMSO and PBS 0.5% BSA (Roche #10735086001)). Caspase activity was assessed 8 hours post treatment according to the manufacturers protocol (Promega #G8091, #G8201) using the PheraStarFSX (BMG Labtech). Data was normalized to the control.

### Genetic depletion by siRNA

HCT-116 cells were seeded at a density of 2×10^6^ cells per 10cm^2^ petri dish. Reverse transfection was performed according to the siTools Biotech protocol using 3nM final concentration of siPOOL control and siPOOL Caspase 8 (siTOOLs Biotech). After 24 hours cells were collected and seeded at 1.5×10^3^ cells/well in a 96-well plate and treated with 15ng/mL recombinant human TNF, 2.5μM RG6146 or control and cell growth was assessed using the IncuCyte S3 Live-cell Analysis System (Essen BioScience) for 7days by acquiring 4 images per well. The remaining cells were re-plated, treated with 15ng/mL TNF for 6 hours and knockdown efficacy was visualized by Western Blot.

### Protein overexpression by viral transduction

Murine colon adenocarcinoma cell line MC38 cells were infected with retroviral murine stem cell virus (MSCV) constructs expressing GFP (MSCV-GFP), murine Bcl-2 and GFP (MSCV-Bcl2-GFP), and murine cFLIP and GFP (MSCV-cFLIP-GFP) then sorted based on GFP expression, as previously described (*47*). MKN45 cells were infected using lentivirus expressing *BIRC2* (GeneID: 321827), *BIRC3* (GeneID: 321828 or for double overexpression GeneID: 327673) or control cDNA (GeneID: 321830) provided by Atum (pD2107-CMV: CMV-ORF, Lenti-ElecD).

### Western blot analysis

Cells were plated at a density of 0.15-0.25×10^6^cells/well in a 6-well plate. After 24 hours cells were treated with small molecules and/or recombinant human TNF or control. After 2-48 hours, cells were collected using 75μl Lysis Buffer (Cell Signaling Technology #9803, Phosphatase Inhibitor Cocktail Set II #524625 and Protease Inhibitor Cocktail Set III #539134 both Calbiochem). Upon cell lysis using a sonicator, protein concentration was assessed with the DC Protein Assay (BioRad #5000112), normalized and Laemmli Buffer (AlfaAeser #J61337) added. Membranes were probed with 1:1000 diluted Anti-cIAP1 (Abcam #ab108361), Anti-cIAP2 (#3130), Anti-TRAF1 (#4715), Anti-Caspase-8 (#4790), Anti-cleaved PARP (#5625), 1:6000 diluted Anti-Actin (#4970) and 1:50000 diluted Anti-Vinculin (#13901) all Cell Signalling Technology. Secondary Goat-anti-rabbit Antibody (Jackson Immuno Research #111-035-144) was diluted 1:5000 and protein bands were visualized with Western Bright Quantum and Sirius HRP Substrate (Advansta) and the Fusion FX (Vilber Lourmat). For analysis of MC38 cells, cells were washed once in ice-cold PBS prior to whole cell lysis using Lamelli buffer (60mM Tris HCl pH 6.8, 10% v/v glycerol, 2% v/v glycerol SDS) and incubated at 95°C for 5-10 minutes. Cell lysate protein concentration was measured using Pierce BCA Protein Assay Kit (ThermoFisher Scientific, 23225) according to the manufacturer’s instructions. Prior to running SDS-PAGE, protein lysates were prepared with sample loading buffer (120mM Tris HCl pH 6.8, 20% v/v glycerol, 4% w/v SDS, 71.5mM β-mercaptoethanol, bromophenol blue). Protein lysates were separated on Mini-PROTEAN TGX 4-15% gels (Bio-Rad, 465-1086) prior to transfer at 0.25A onto Immobilon-P (IPVH00010) membranes in transfer buffer (25mM Tris HCl, 192mM Glycine, 5% v/v methanol) at 4°C. Next, membranes were incubated overnight using the following primary antibodies: anti-PARP (Cell Signaling, 46D11), anti-Caspase-3 (Cell Signaling, D3R6Y), and Actin (Sigma, A2228). Membranes were incubated with horseradish peroxidase (HRP)-conjugated secondary antibodies at room temperature for 1 hour and washed at least 3 times in TBS supplemented with 0.1% v/v Tween20. Immunoreactive bands were detected using ECL reagents (Amersham ECL or ECL Prime, GE Healthcare) by film exposure (Fujifilm Super RX, Fujifilm) using an Agfa CP1000 developer (Agfa).

### RNA-sequencing of HCT-116 and MKN45 cells

Cells were plated at a density of 0.25×10^6^ cells/well in a 6-well plate. After 24 hours cells were treated with 2.5μM RG6146 and 20ng/mL recombinant human TNF or control. Cell were harvested after 2 hours and frozen in liquid nitroge. RNA was extracted using the Qiagen RNeasy Mini Kit (#74104), as per manufacturer’s instructions except that lysate was loaded on a QIAshredder column before loading on a spin column. Sequencing libraries were generated from 100ng input RNA using the Illumina TruSeq Stranded mRNA LT Sample Preparation Kit (Set B, #RS-122-2102) as per manufacturer’s instruction. Libraries were sequenced on an Illumina HiSeq4000 using paired-end sequencing 2×50bp reads to an average depth of 18 to 37 million sequences per sample. Base calling was performed with BCL to FASTQ file converter bcl2fastq (v2.17.1.14) from Illumina (https://support.illumina.com/downloads.html). In order to estimate gene expression levels, paired-end RNASeq reads were mapped to the human genome (hg38) with STAR aligner (v2.5.2a) using default mapping parameters (*48*). Numbers of mapped reads for all RefSeq transcript variants of a gene (counts) were combined into a single value by using SAMTOOLS software (*49*).

### 3’-mRNA sequencing of MC38 and E0771 cells

MC38 and E0771 cells were treated as indicated prior to immediate RNA extraction by resuspension in TRIzol reagent (Thermo Fisher Scientific) and isolation using the Direct-Zol RNA Miniprep kit (Zymo Research), as per manufacturer’s instructions. Sequencing libraries were generated from 500ng input RNA using the QuantSeq 3’RNA-seq Library Prep Kit for Illumina (Lexogen), as per manufacturer’s instructions. Libraries were sequenced on an Illumina NextSeq 500 with single-end 75bp reads to a depth of 5M reads per sample. Raw sequencing reads were demultiplexed with bcl2fastq (v2.17.1.14) and subsequently trimmed at the 5’-end and 3’ ends (to remove poly-A-tail derived reads) using CutAdapt (v1.14). Reads were then mapped to reference genomes (mm10) using HISAT2 (v2.1.0) software. Counting of reads across annotated genomic features (exons) was conducted using featureCounts (subread package, v1.5.2). Differential gene expression analysis was performed using the VoomLimma workflow to determine statistical significance. All further RNA-seq data analysis and figures generation was performed using Rstudio (v3.6.1) software. GO Term analysis was performed using ToppGene software. Gene Set Enrichment Analysis (GSEA) software (v3.0) was used for identification of enriched gene sets, using the MSigDB KEGG and Hallmarks datasets. Human gene expression array data on human umbilical vein endothelial cells (HUVEC) treated with recombinant TNF in the presence or absence of JQ1 was downloaded from Gene Expression Omnibus (GEO; accession number: GSE53999) and analysed by Limma in Rstudio.

### Quantitative RT-PCR

Cells were plated at a density of 0.25×10^6^ cells/well in a 6-well plate. After 24 hours, cells were treated with RG6146 and/or recombinant human TNF or control. Cells were harvested after 2 hours of treatment and RNA was isolated with the RNeasy Mini Kit (Qiagen #74104). RNA levels were assessed using qScript™ XLT One-Step RT-qPCR ToughMix® (Quanta Bioscience #95132-100) and the corresponding TaqMan Gene Expression Assays (Thermo Fisher #Hs02786624_g1 (GAPDH), #Hs01112284_m1 *(BIRC2*), #Hs00985031_g1 (*BIRC3*), #Hs01090170_m1 (*TRAF1*)) using the LightCycler® 480 System (Roche). The 2ΔΔCt-Method was used to calculate the relative RNA expression and normalized to Actin or GAPDH housekeeping genes.

### Viability Assay

HCT-116 and MKN45 cells, and genetically modified variants thereof, were plated at a density of 0.005×10^6^ cells/well in a 96-well plate. After 24 hours, cells were treated with a dose response of compound and recombinant human TNF or control. Viability was measured after 72 hours using 50μL CTG2.0 and the PheraStarFSX.

### Generation of TNFR1 and TNFR2 knockout cell lines

Recombinant *S. pyogenes* Cas9 nuclease (Alt-R® S.p. Cas9 Nuclease V3) was ordered from IDT and the following synthetic guide RNA (gRNA) were ordered from Synthego in 1.5nM yield:

Tnfsf1a sgRNA: AGCAGAGCCAGGAGCACCUG Tnfsf1b

sgRNA: UACCCAGGUUCCGGUUUGUA

gRNAs were re-suspended in TE buffer at 200µM and stored at -20ºC. CRISPR ribonucleoprotein (RNP) complexes were made by mixing 100pmol Cas9 and 1.5µL reconstituted gRNA in sterile H_2_O (total volume 5µL) and incubated at room temperature for 20 minutes. In the meantime, MC38 cells were trypsinized, washed twice with PBS and 0.25×10^6^ cells per condition resuspended in 20μL supplemented cell line solution (SF Cell Line 4D-NucleofectorTM X Kit S, Lonza). Cells were then transferred to the RNP solution, gently mixed, and transferred to the well of a 16-well Nucleocuvette™ strip (SF Cell Line 4D-NucleofectorTM X Kit S, Lonza). Cells were electroporated on a 4D-NucleofectorTM X Unit (Lonza) following the manufactuers reccomendations for HEK-293T cells (pule code: CM-130). 100µL of pre-warmed media was transferred into each well, mixed gently, then transferred to into a pre-warmed 6-well plate containing 2mL of media. Cells were expanded and those deficient in TNFR1 and/or TNFR2 were subsequently selected by multiple rounds of FACS sorting.

### Assay for Transposase Accessible Chromatin with high-throughput sequencing (ATAC-seq)

Assay for Transposase Accessible Chromatin using Sequencing (ATAC-Seq) was performed using an improved protocol to reduce mitochondria from the transposition reaction. Briefly, 5×10^5^ MC38 cells were treated as indicated with recombinant murine TNF (10ng/mL) in the presence or absence of RG6146 (2.5µM). Cells were washed once in ice-cold PBS and lysed in ATAC lysis buffer (0.1% Tween-20, 0.1% NP-40, 3mM MgCl_2_, 10mM NaCl, 10mM Tris HCl pH 7.4). Tagmentation was performed with Tn5 transposase and 2x TD Buffer (Nextera DNA Library Prep Kit, Ilumina) for 30 minutes at 37°C. Tagmented DNA was purified using a MinElute colum (Qiagen, 28004) and amplified for 12 cycles using 2x KAPA HiFi HotStart ReadyMix (Kapa Biosystems, KK2602). The amplified libraries were purified using MinElute columns (Qiagen) and sequenced on an Ilumina NextSeq 500 with 75 bp single-end reads. Library QC and quantification was performed using D1000 high-sensitivity screen tape with 4200 TapeStation Instrument (Agilent Technologies) and size selected for between 200 bp and 500 bp using a Pippin Prep system (Sage Science).

### Chromatin immunoprecipitation and sequencing (ChIP-seq)

MC38-Ova cells were treated as indicated for 3 hours prior to aspiration of media and one wash in ice-cold PBS. Cells were crosslinked directly in tissue culture plates with 1% formaldehyde for 20 minutes at room temperature and subsequently quenched with 1.25M glycine. Cells were manually scraped from plates, collected, and washed three times with ice-cold nuclear extraction buffer, prior to lysis in ChIP lysis buffer (20 mM Tris-HCl, 150 mM NaCl2, 2 mM EDTA, 1% IGEPAL CA-630 (Sigma-Aldrich) and 0.3% SDS in water). Sonication was performed using a Covaris ultrasonicator at maximum power for 16 minutes to achieve an average DNA fragment size of 300-500bp. Immunoprecipitation reactions were performed overnight at 4ºC by diluting sonicated chromatin 1:1 in ChIP dilution buffer (20 mM Tris-HCl, 150 mM NaCl2, 2 mM EDTA, 1% Triton-X, and phosphatase/protease inhibitors in water) and addition of a 1:1 ratio of protein A and G magnetic beads (Life Technologies) were used to bind crosslinked protein/DNA. The following antibodies were used for ChIP-seq: NF-KB p65 polyclonal antibody (Diagenode, C15310256), H3K27Ac (Abcam, ab4729), and BRD4 (Bethyl, A301-985A100). Samples were washed once with ChIP dilution buffer, wash buffer 1 (0.1% SDS, 1% Triton X-100, 2mM EDTA, 500mM NaCl, 20mM Tris-HCl pH 8), wash buffer 2 (0.5% deoxycholate, 0.5% NP-40, 2mM EDTA, 250mM LiCl, 20mM Tris-HCl pH 8), and TE buffer (1mM EDTA, 10mM Tris-HCl pH 7.5) prior to incubation in reverse crosslinking buffer (200mM NaCl, 100mM NaHCO_3_, 1% SDS, 300μg/mL Proteinase-K) for 4 hours at 55°C with shaking. Finally, the supernatant was reverse-crosslinked overnight (12-16 hours) at 65°C prior to ChIP DNA isolation using Zymogen ChIP DNA Clean and Concentrator Kit (Zymo Research, D5205). For ChIP-seq, libraries were generated using the NEBNext Ultra II DNA Library Prep Kit (NEB, E7645) and sequenced on an Ilumina NextSeq 550 with 75 bp single-end reads. Library QC and quantification was performed using D1000 high-sensitivity screen tape with 4200 TapeStation Instrument (Agilent Technologies) and size selected for between 200 bp and 500 bp using a Pippin Prep system (Sage Science).

### Analysis of ChIP-seq and ATAC-seq datasets

For analysis of both ChIP-seq and ATAC-seq datasets, raw sequencing reads were demultiplexed using bcl2fastq (v2.17.1.14) and quality control performed with FASTQC (v0.11.5). Adaptor sequences were removed and reads aligned to the human reference genome (mm10) using CutAdapt (v1.14) and Bowtie2 (v2.3.5), respectively. Samtools (v1.4.1) was used for processing of SAM and BAM files, from which Model-based Analysis of ChIP-Seq (MACS2, v2.2.5) peak calling was performed. BAM files were converted into BigWig files using the bamCoverage function (Deeptools, v3.0.0) using the following settings (--normalizeUsing CPM --smoothLength 150 --binSize 50 -e 200 scaleFactor 1). BigWig files were imported into Integrative Genomics Viewer (IGV, v2.7.0) for visualization of specific loci. Using Deeptools (v3.0.0), average profile plots and heatmaps were generated by computing read average read density (from BigWig files) across defined genomic intervals using computeMatrix, which we subsequently plot using plotProfile and plotHeatmap, respectively. Genomic regions spanning mm10 blacklist intervals (ENCODE) were excluded using Deeptools. Annotation of putative super enhancer regions from H3K27ac ChIP-seq data was performed using Ranking Ordering of Super Enhancer (ROSE, v1.0.5) using a 12.5 k.b. stitching distance and a 2.5 k.b. TSS exclusion to reduce promoter bias. Peak calling was performed with MACS2 with default parameters. Subsequent annotation of ATAC-Seq/ChIP-Seq peaks to proximal genes and motif analyses was performed using annotatePeaks.pl and findPeaks (Homer, v4.8).

### CEA-TCB Assay

MKN45 cells were plated at a cell density of 0.025×10^6^ cells/well in a 96-well plate and frozen PBMCs were thawed and kept in the incubator overnight in RPMI + GlutaMax + 10% hiFBS. 0.25×10^6^ PBMCs/well and CEA-TCB (Roche) were added to the co-culture. 48-72 hours post treatment, cells were centrifuged and cell killing assessed using the LDH Assay (Roche #11644793001) according to the manufacturers protocol using the SPECTROstar Nano (BMG Labtech).

### Assessment of co-culture supernatant cytotoxicity

Co-culture was prepared as described for the CEA-TCB assay using 20nM CEA-TCB. Supernatant was collected after 24 hours and filtered through a 0.22μM Filter. HCT-116 and MKN45 cells were plated the day before at a density of 0.005×10^6^ cells/well in a 96-well plate, medium was removed and supernatant of the co-culture was added on the cells. Cells were treated with a dose response of RG6146 or the corresponding control. Where indicated 10μg/mL TNF-alpha Monoclonal Antibody or ITC were added. Cell viability was measured by adding 50μL CTG2.0 after 72 hours and the PheraStarFSX.

### Tumor antigen-independent bystander killing assay

HCT-116-GFP cells and MKN45-RFP cells, expressing GFP and RFP, respectively, were plated at a cell density of 0.02×10^6^ cells/well in 96-well plates. CEA-TCB (40nM) and 0.2×10^6^ PBMCs/well were added after 24 hours to the co-culture. RG6146 or control were added and pictures were taken by the Incucyte S3 Live-Cell Analysis System (Essen Bioscience) for 72 hours (4 images/well). Data was analyzed using the Incucyte S3 Live-Cell Analysis System looking at the GFP count per image.

### *In vivo* assessment of CEA-TCB and JQ1 anti-tumor activity

The MC38_HOMSA_CEACAM5 transfected cell line was cultured in DMEM high-glucose medium supplemented with NEAA, 4 mM glutamine, 2 mM sodium pyruvate, 10% FBS, 500 µg/mL G-418 at 37 °C and 5 % CO_2_. MC38-CEA cell were injected subcutaneously (sc) at a concentration of 5×10^5^ together with matrigel into female C57/Bl6 huCEA tg mice (Charles River, France), aged 5-8 weeks at arrival. Twice weekly tumor volume was measured in two dimensions (a = length; b = width) with a caliper. Tumor volume (V) was determined by the following equation: V = ab2/2. Animal treatment started after randomization when mean tumor size was about 100 or 130mm^3^ (10 mice / group). CEA-TCB antibody was administered at 2.5mg/kg iv twice weekly (4x). 50mg/kg JQ1 (HY-13030, MedChemExpress) was given intraperitoneal (i.p.) once daily (14x). 2mg/kg anti-TNF-α Mab (BioLegend, #506347) was injected intravenous (i.v.) twice weekly (4x). Tumor growth inhibition (TGI) for each group was calculated according to the following formula: (1 - [T - To] / [C - Co]) x 100 where T= Mean tumor volume of mice in the same treatment group (last measurement), To= Mean tumor volume of mice in the same treatment group (first measurement), C= Mean tumor volume of mice in the control group (last measurement), Co= Mean tumor volume of mice in the control group (first measurement). The %-change in tumor volume for each mouse was calculated with the following equation: ([M - Mo] / [Mo]) x 100 where M= Tumor volume (last measurement) and Mo= Tumor volume (first measurement).

### *In vivo* assessment of anti-PD1 and JQ1 anti-tumor activity

Male 6-8 week old C57BL/6 mice were injected with MC38 cells sc and randomized 6 days post tumor inoculation into four groups containing 10-13 animals/ group. Mice were treated daily with 50mg/kg JQ1 or DMSO control i.p. (days 6-22 post tumor inoculation) and with 0.2mg/kg anti-PD1 (clone RMP1-14) or isotype control (ITC; clone 2A3) i.p. on days 9, 13, 26 and 20 post tumor inoculation. Twice a week, tumor volume was measured in two dimensions (a = length; b = width) with a caliper. Tumor volume (V) was determined by the following equation: V = ab2 / 2. Results contain data from two biological independent experiments.

### Chemical Structure of RG6146

The chemical structure of RG6146 in Figure 8 was generated with ACD/ChemSketch (v2019.2.2, Advanced Chemistry Development, Inc., Toronto, ON, Canada)

### Intracellular cytokine measurement of OTI T-cells

Activated OTI T-cells were combined with MC38-OVA tumor cells at a 1:4 effector:target ratio and cultured together with 0.6μL/mL GolgiStop (BD Biosciences) for 5 hours. Co-cultures were harvested and then stained with fixed viability dye (Zombie Aqua, Biolegend) and fluorochrome-labelled T-cell-specific antibodies targeting CD5 (clone 53-7.3, BD Biosciences) and CD44 (clone IM7, Biolegend). Labelled cells were fixed with 4% paraformaldehyde for 10 minutes, washed with staining buffer, resuspended in Perm/Wash buffer (BD Biosciences) for 15 minutes and then stained with fluorochrome-labelled antibodies targeting TNFα and IFNγ (clones MP6-XT22 and XMG1.2, eBioscience). Labeled cells were subsequently examined via flow cytometry (BD LSR Fortessa X-20, BD Biosciences).

### Cytometric bead array (CBA)

Detection of secreted cytokines in co-culture supernatants was performed using a mouse (BD Biosciences, 552364) or human inflammation CBA kit (BD Bioscience #558264, #560112, #560304, #558269) as per manufacturer’s instructions. Beads detecting cytokines from mouse were analyzed on a FACS Verse (BD Biosciences, North Ryde, New South Wales, Australia). All assays were analyzed using triplicate determinations. Beads detecting cytokines from human were analyzed on an IQue Screener Plus (Sartorius). All assays were analyzed using quintuplicate determinations.

### Assessment of CEA expression by Flow Cytometry

CEA-level on HCT-116-GFP and MKN45-RFP cells was assessed by plating 0.1×10^6^ cells/well in a 6-well plate. Cells were stained with 1:500 diluted Zombie NIR (BioLegend #423105) for 20 minutes and washed with PBS + 0.5% BSA + 2mM EDTA. 10μg/mL CEA-Antibody (Novus BioScience #NBP2-34594-0.1 mg) or Isotype Control (mo IgG1 Isotype, BioLegend #401402) were added to the cells for 30minutes. The secondary antibody (Southern Biotech #1010-31) was diluted 1:100 and added for 30minutes. CEA level was assessed by the Cytoflex S Benchtop FACS. Data was analyzed using FlowJo_V10.

### Bystander Killing Chromium Release Assay

Bystander killing activity was measured using a standard chromium release assay as previously described *12*. Briefly, MC38 cells were labeled with 100 μCi of ^51^Cr (Perkin Elmer) and mixed 50:50 with non-^51^Cr-labelled MC38-Ova cells. Activated OTI T-cells were then added to the targets at the indicated E:T ratios in the presence of RG6146 (2.5μM) or DMSO vehicle. After 18 hours of incubation (37 °C, 10% CO2), co-culture supernatants were harvested, and the level of ^51^Cr was quantified by a Gamma counter (Wallac Wizard). Percentage specific killing was determined using the formula: (Sample 51Cr release – Spontaneous background ^51^Cr release)/ (Total ^51^Cr release – Spontaneous Background ^51^Cr release) × 100%, and represented as a Michaelis–Menten kinetic trend. All assays were performed using technical triplicates.

---

**Supplementary Video 1. Time-lapse microscopy of MC38-Ova cells co-culutred with perforin-deficient OTI T-cells in the presence of DMSO**

A representative field of view from time-lapse microscopy performed to visualize MC38-Ova cells co-cultured with a fixed ratio of perforin-deficient OTI T-cells pre-labelled with cell trace violet (appearing blue in video), and cultured for 24 hours in the presence of DMSO vehicle and a fluorescent active caspase-3/7 reporter (appearing as green ‘flashes’ in video). Time scale indicates hours:minutes since addition of OTI T-cells to the co-culture supernatant.

**Supplementary Video 2. Time-lapse microscopy of MC38-Ova cells co-culutred with perforin-deficient OTI T-cells in the presence of RG6146**

A representative field of view from time-lapse microscopy performed to visualize MC38-Ova cells co-cultured with a fixed ratio of perforin-deficient OTI T-cells pre-labelled with cell trace violet (appearing blue in video), and cultured for 24 hours in the presence of 2.5µM RG6146 and a fluorescent active caspase-3/7 reporter (appearing as green ‘flashes’ in video). Time scale indicates hours:minutes since addition of OTI T-cells to the co-culture supernatant.

**Supplementary Video 3. Time-lapse microscopy of HCT-116 to show Caspase-8 depletion partially rescuses the combination effects of RG6146 and TNF**

Representative field of view from time-lapse microscopy performed to visualize growth of HCT-116 cells depleted of Caspase 8 or control in the presence of RG6146 and TNF. HCT-116 cells were reverse transfected with a scrambled control prior to treatment DMSO (left quarrant) or treated with 15 ng/mL recombinant human TNF and 2.5 μM RG6146 (second from left quadrant) for 6 days and 20 hours. HCT-116 cells were also reverse transfected with siPOOLs targeting Caspase-8 (siCasp8) and treated with DMSO (second from right quadrant) or treated with 15 ng/mL recombinant human TNF and 2.5 μM RG6146 (right quadrant) for a duration of 6 days and 20 hours.

**Supplementary Figure 1.**
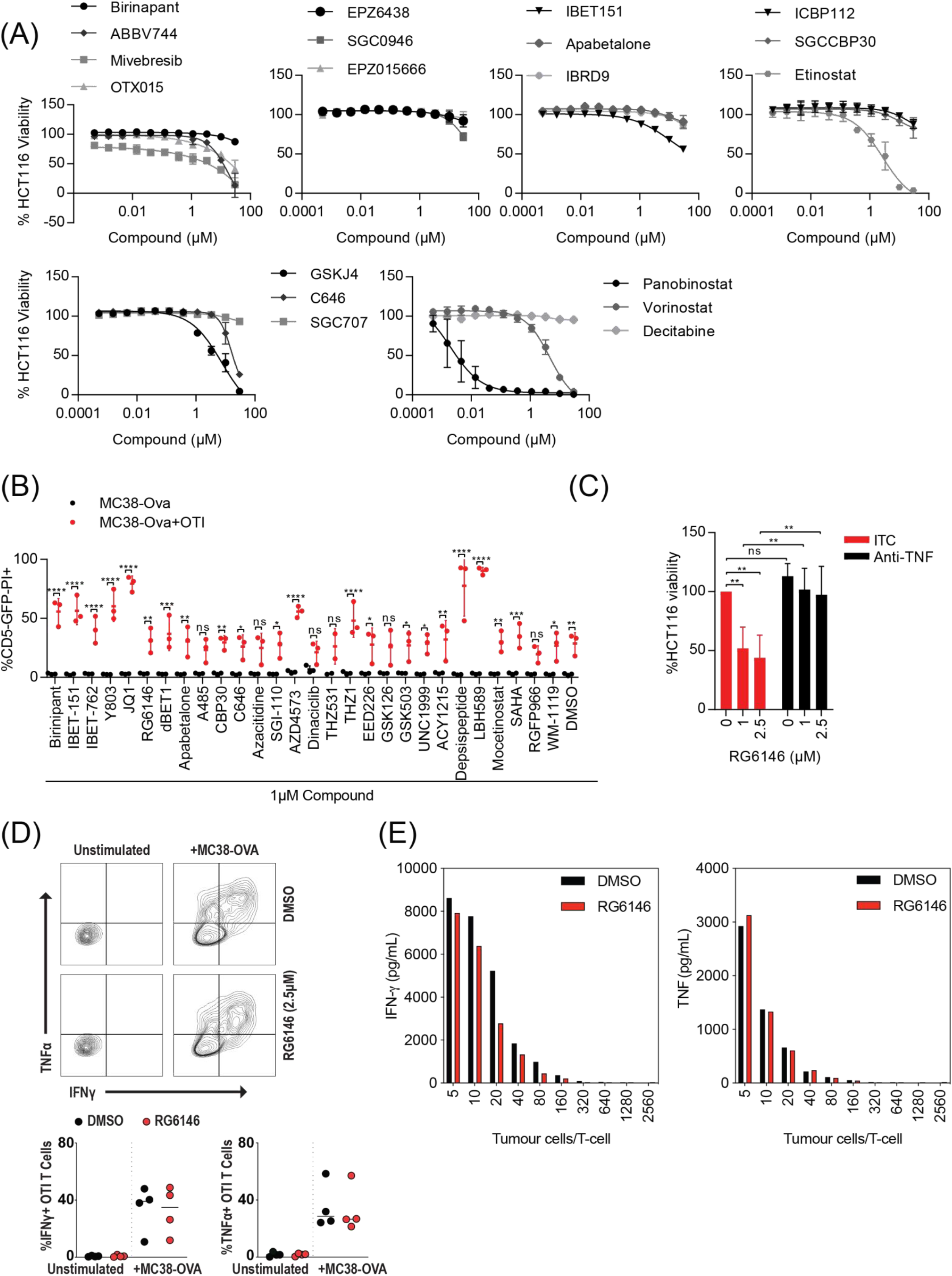
BET Bromodomain inhibition augments CD8^+^ T-cell anti-tumor immunity. **(A)** HCT-116 cells were treated with increasing doses of agents from a bespoke small molecule library of epigenetic therapies, as well as SMAC mimetics. Viability (%) was assessed following 72 hour drug treatment and data was normalized to DMSO control. Data represents mean of three independent experiments. **(B)** MC38-Ova^+^GFP^+^ cells were treated with small molecule epigenetic therapies (1µM) or DMSO in the presence or absence of OTI T-cells (4:1 tumor cells/OTI) for 13-14 hours prior to assessment of tumor cell death by flow cytometry (PI^+^GFP^-^CD5^-^). Data represents mean of three independent experiments. Statistical significance denotes difference between MC38 mono and co-culture to treatment (2-way ANOVA, *p<0.05, **p<0.01, ***p<0.001, ****p<0.0001). **(C)** NLV peptide loaded on HCT-116 cells co-cultured with CMV-specific T cells were treated with 1-2.5μM RG6146 or DMSO control for 48 hours in the presence of TNF neutralizing antibody (Anti-TNF) or isotype control (ITC). Cell viability was normalized to DMSO and data represents mean of three independent experiments. Statistical significance denotes difference between DMSO and RG6146 as well as Anti-TNF and ITC (2-way ANOVA, **p<0.01). **(D)** Intracellular cytokine staining in OTI CD8^+^ T-cells alone (unstimulated) or co-cultured with MC38-Ova cells and treated with RG6146 (2.5µM) or DMSO vehicle for 5 hours in the presence of GolgiStop prior to intracellular flow cytometry staining of CD5^+^CD44^+^ T cells for TNF and IFN-y. Data represents mean of four independent experiments. **(E)** Cytokine secretion detected in MC38-Ova and OTI co-cultures treated with RG6146 (2.5µM) or DMSO vehicle for 18 hours prior to collection of culture medium and analysis of TNF and IFN-y cytometric bead array from a representative experiment.

**Supplementary figure 2.**
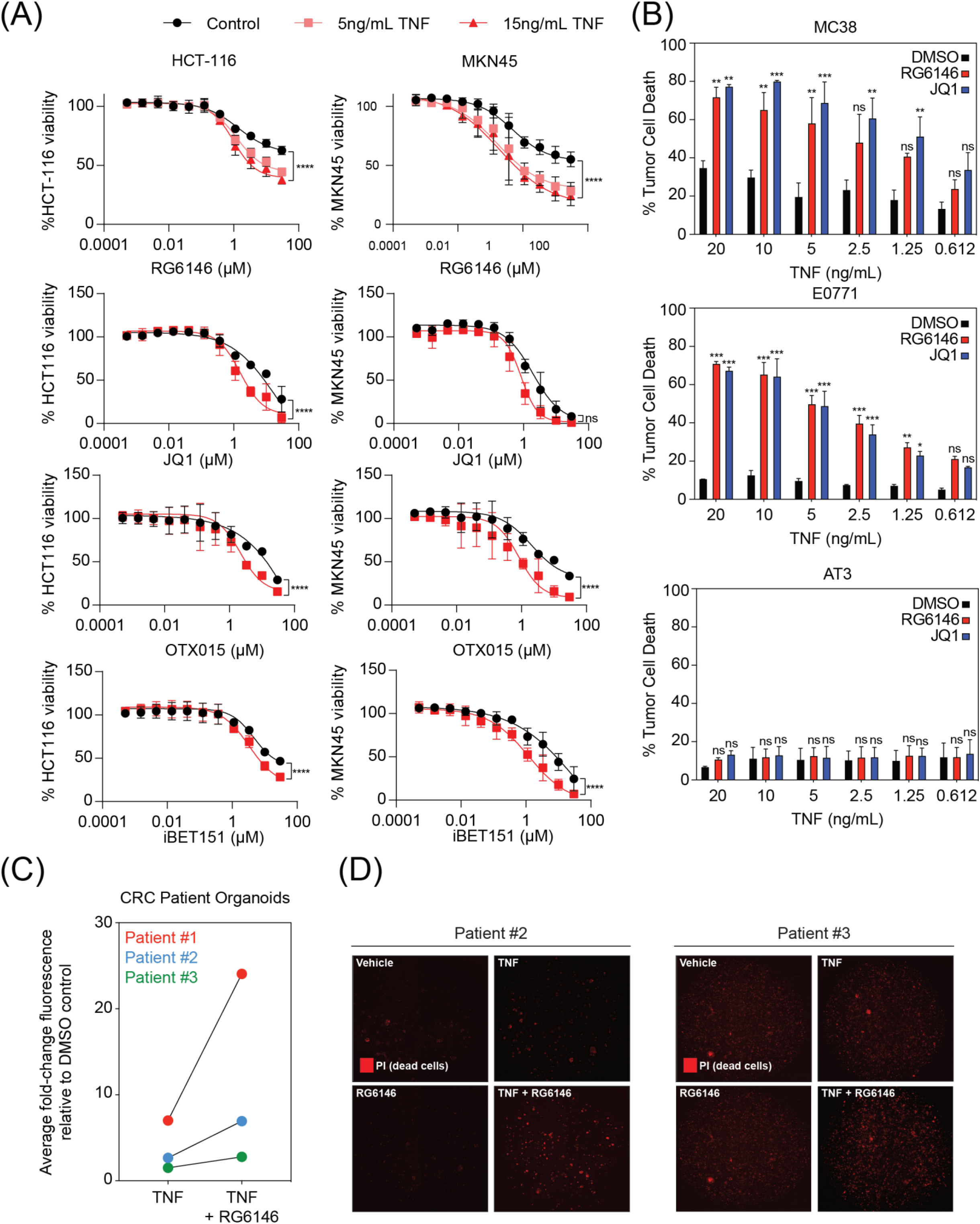
BET inhibition potentiates the cytotoxic effects of TNF. **(A)** HCT-116 and MKN45 cells were treated for 72 hours with increasing concentrations of RG6146, JQ1, OTX015 or iBET151 in the presence or absence of 5 and 15ng/mL recombinant human TNF prior to assessment of tumor cell viability (%). Viability was normalized to DMSO control and data represents mean of three independent experiments. Statistical significance denotes difference between highest concentration of RG6146 single agent and combination with 15ng/mL TNF (2-way ANOVA, ****p<0.0001). **(B)** Murine solid tumors MC38, E0771, and AT3 were treated for 24 hours with increasing concentrations of recombinant murine TNF in the presence of JQ1 (1µM), RG6146 (2.5µM), or DMSO control prior to assessment of tumor cell death by flow cytometry (propidium iodide positivity). Data represents mean of three independent experiments. Statistical significance denotes difference between DMSO and JQ1/RG6146 at each ratio (2-way ANOVA, *p<0.05, **p<0.01, ***p<0.001). **(C)** Log fold-change in PI fluorescence in organoids derived from three independent CRC patient specimens treated with TNF (10ng/mL) or TNF+RG6146 (2.5µM) for 24 hours. **(D)** Fluorescence microscopy of two independent patient-derived CRC organoid cultures and exposed to combinations of RG6146 (2.5µM) or DMSO and recombinant TNF (10ng/mL) in media containing propidium iodide (PI).

**Supplementary figure 3.**
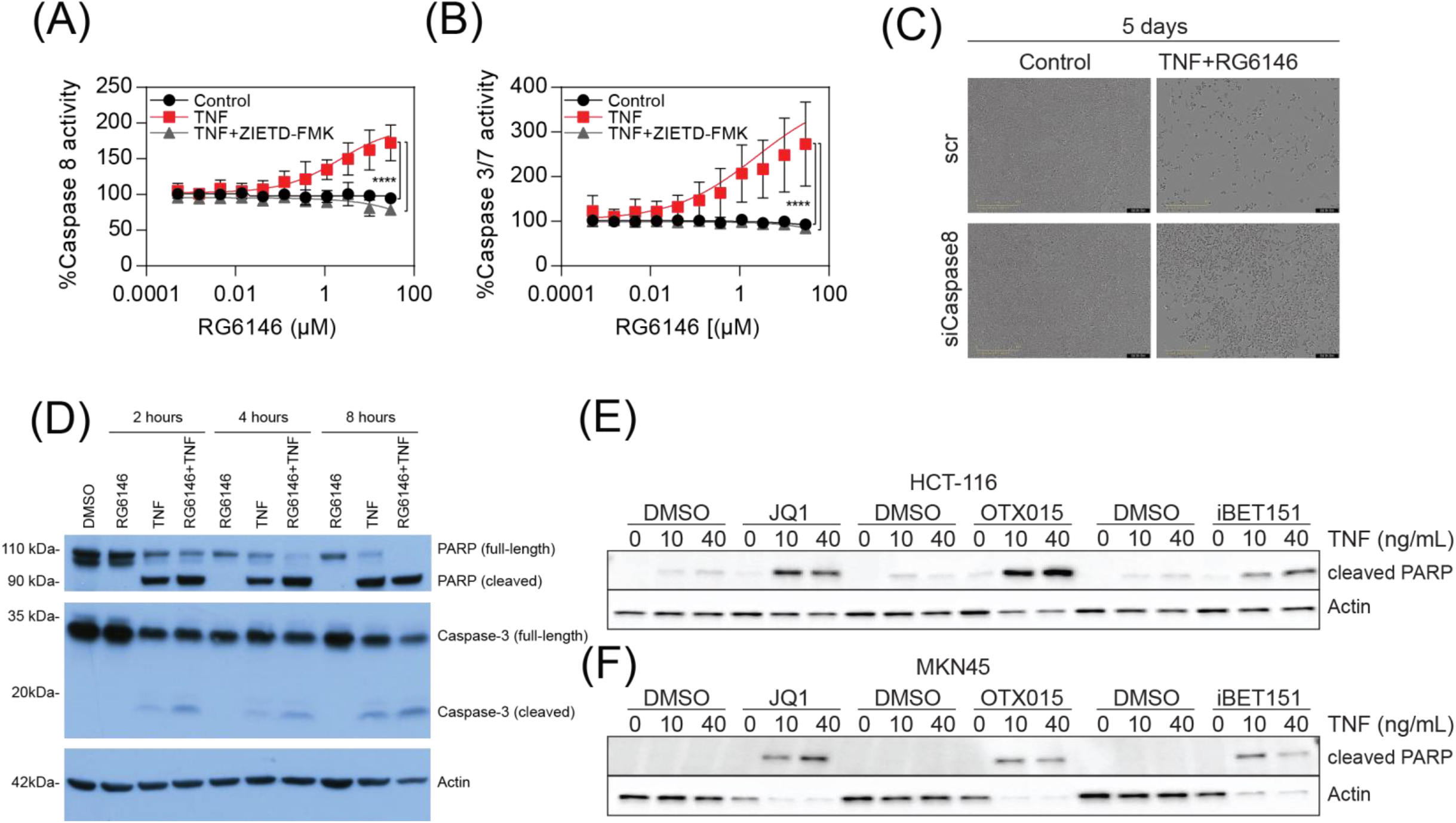
Inhibition of BET proteins enhances caspase activity following TNF stimulation. Analysis of **(A)** Caspase-8 and **(B)** Caspase-3/7 activity in MKN45 cells by Caspase-Glo following 8 hours treatment with increasing concentrations of RG6146 in the presence of 15 ng/mL recombinant human TNF and 1μM Caspase-8 inhibitor (ZIETD-FMK). Data is normalized to each DMSO control and represents mean of three independent experiments. Statistical significance denotes difference between highest concentration of RG6146 single agent vs combination with TNF +/- ZIEDT-FMK (2-way ANOVA, ****p<0.0001). **(C)** HCT-116 cells were reverse transfected with siPOOLs targeting Caspase-8 or scramble control. Cells were seeded the next day and treated with 15ng/mL recombinant human TNF and 2.5μM RG6146 or control. Representative images of cell confluence after 5 days are shown. **(D)** Immunoblot analysis of PARP and Caspase-3 protein in MC38 cells following treatment with recombinant murine TNF (10ng/mL), RG6146 (2.5µM), or the combination for 2, 4, and 8 hours. Actin was used as a loading control. Data is one representative blot of three independent experiments performed. (**E-F**) Immunoblot analysis of PARP protein from **(E)** HCT-116 or **(F)** MKN45 cells treated with 2.5μM JQ1, OTX015 or iBET151 and 10 or 40ng/mL recombinant human TNF for 24 hours. Actin was used as a loading control. Data is one representative blot of three independent experiments performed.

**Supplementary figure 4.**
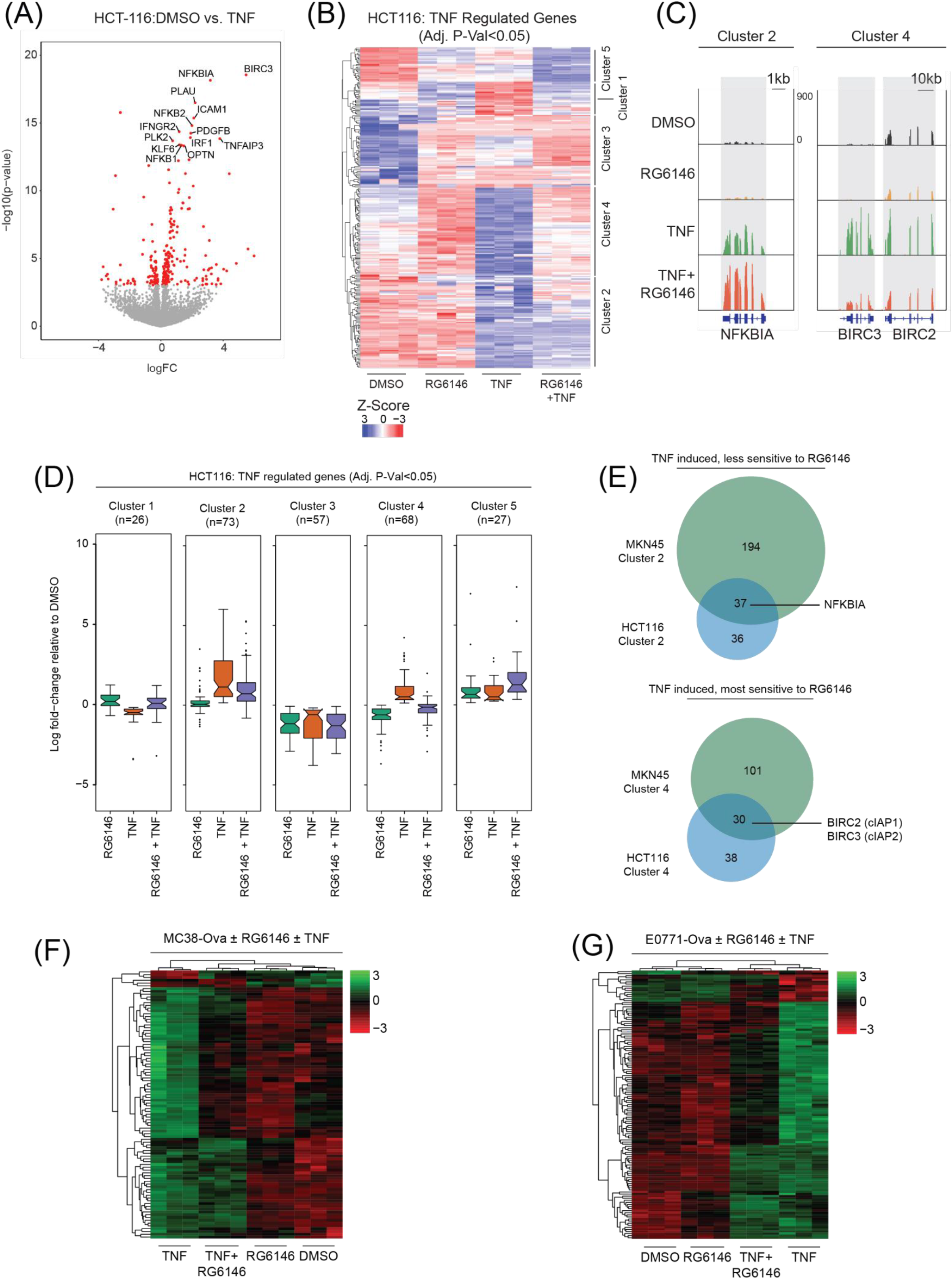
BET inhibition modulates transcription downstream of TNF stimulation. **(A)** RNA-seq from HCT-116 cells treated in triplicate with recombinant human TNF (20 ng/mL) or DMSO vehicle. Differentially expressed genes (red) are defined as adjusted P-value<0.001. **(B)** Heatmap of normalized log CPM of genes differentially expressed in response to TNF (adj. P-value<0.001) separated into 5 groups by k-means clustering. **(C)** IGV genome browser screenshot showing RNA-seq signal at NFKBIA (representative Cluster 2 gene) and BIRC2/3 loci (representative Cluster 4 loci). **(D)** Boxplot of log fold-change values comparing RG6146, TNF, or the combination of RG6146 and TNF, with DMSO for genes within each of the 5 clusters. **(E)** Overlap of genes from MKN45 and HCT-116 RNA-seq data. **(F)** Heatmap of normalized log CPM of genes differentially expressed in response to TNF in MC38-Ova cells treated with RG6146 (2.5µM) alone or in combination with recombinant murine TNF (20ng/mL), or vehicle control. **(G)** Heatmap of normalized log CPM of genes differentially expressed in response to TNF in E0771-Ova cells treated with RG6146 (2.5µM) alone or in combination with recombinant murine TNF (20ng/mL), or vehicle control.

**Supplementary figure 5.**
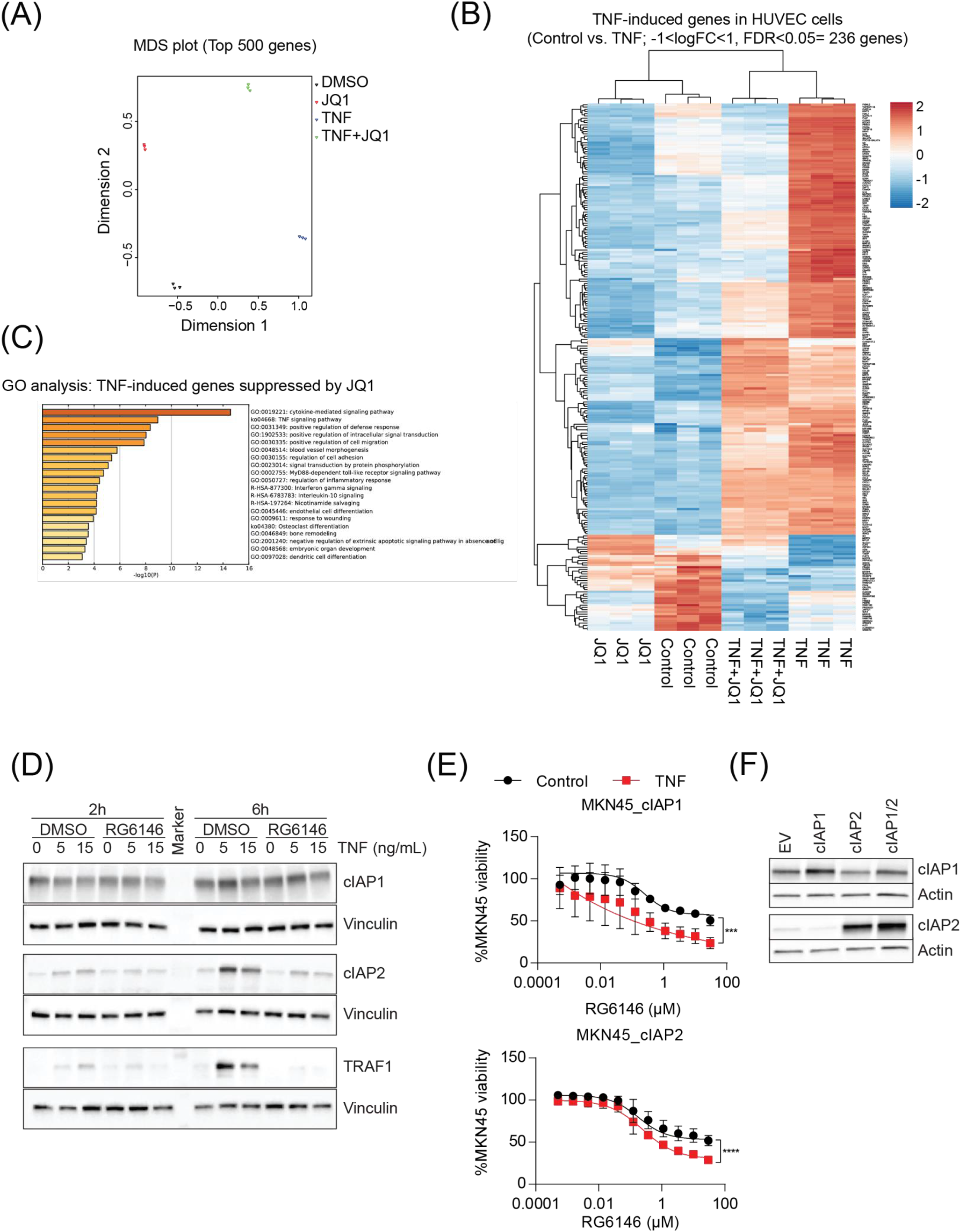
Transcriptional modulation of NF-κB leads to suppression of key negative regulators of TNF. **(A)** Multidimensional scaling (MDS) plot of microarray data (GSE53999) from human endothelial cells treated in triplicate with JQ1 (500nM) in the presence or absence of recombinant human TNF (25ng/mL). **(B)** Heatmap of normalized log counts of genes differentially expressed in response to TNF (adj. P-value<0.05 and -1<log fold-change>1). **(C)** Gene ontology (GO) enrichment of gene signatures in the genes differentially expressed in response to TNF (adj. P-value<0.05 and -1<log fold-change>1). **(D)** Immunoblot of cIAP1, cIAP2, and TRAF1 protein from MKN45 cells treated for 2 and 6 hours with 5 or 15ng/mL recombinant human TNF in the presence or absence of RG6146 (2.5μM). Vinculin was used as a loading control. Representative blot of three independent experiments is shown. **(E)** Viability of MKN45 cells transduced with lentivirus to overexpress cIAP1 (MKN45_cIAP1) or cIAP2 (MKN45_cIAP2) and treated for 72 hours with increasing concentrations of RG6146 and 15ng/mL recombinant human TNF. Data is normalized to DMSO and represents mean of three independent experiments. Statistical significance denotes difference between highest concentration of RG6146 single agent and combination with TNF (2-way ANOVA, ***p<0.001, ****p<0.0001). **(F)** Immunoblot of cIAP1 and cIAP2 protein in MKN45 cells transduced with lentivirus to overexpress cIAP1, cIAP2, or both cIAP1 and cIAP2 (cIAP1/2) or empty vector (EV). Actin was used as a loading control.

**Supplementary figure 6.**
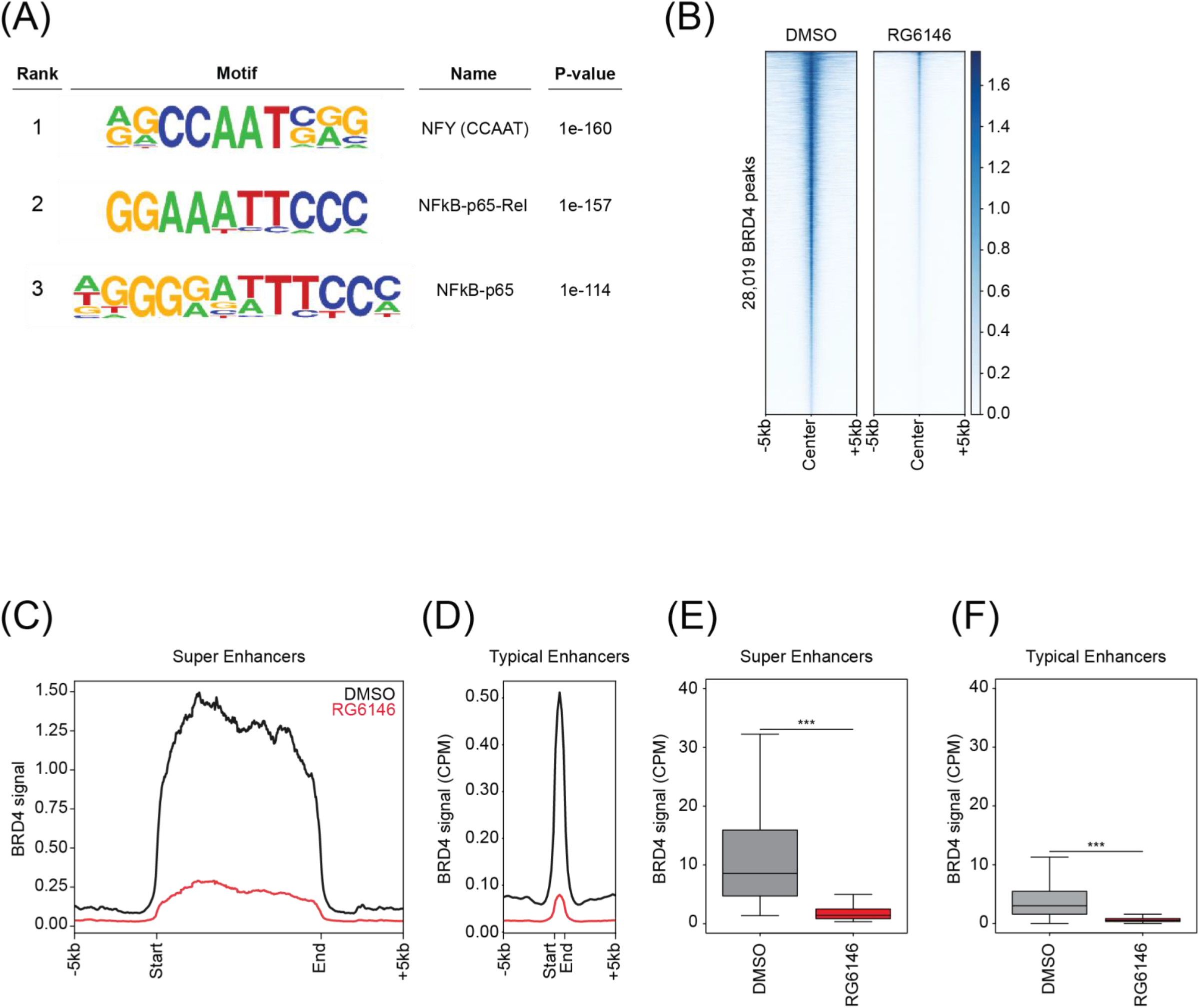
Global eviction of BRD4 from chromatin by RG6146. *De novo* motif analysis of ATAC-seq data from MC38 cells treated for 3 hours with recombinant murine TNF (using ATAC-seq data from control treated MC38 cells as background). Heatmap of normalized BRD4 ChIP-seq signal from MC38 cells treated for 3 hours with RG6146 (2.5µM) or DMSO vehicle (centered on 28,019 BRD4 peaks identified in DMSO vehicle treated cells ± 2.5kb). **(C)** Average profile of normalized BRD4 ChIP-seq signal across H3K27ac-ranked super enhancers in MC38 cells treated for 3 hours with RG6146 (2.5µM) or DMSO vehicle. **(D)** Average profile of normalized BRD4 ChIP-seq signal across typical enhancers (all H3K27ac enhancers excluding those super enhancers) in MC38 cells treated for 3 hours with RG6146 (2.5µM) or DMSO vehicle. **(E)** Boxplot represents quantification of normalized BRD4 ChIP-seq signal at super enhancers (Students t-test, ***p<0.001). **(F)** Boxplot represents quantification of normalized BRD4 ChIP-seq signal at typical enhancers (Students t-test, ***p<0.001).

**Supplementary figure 7.**
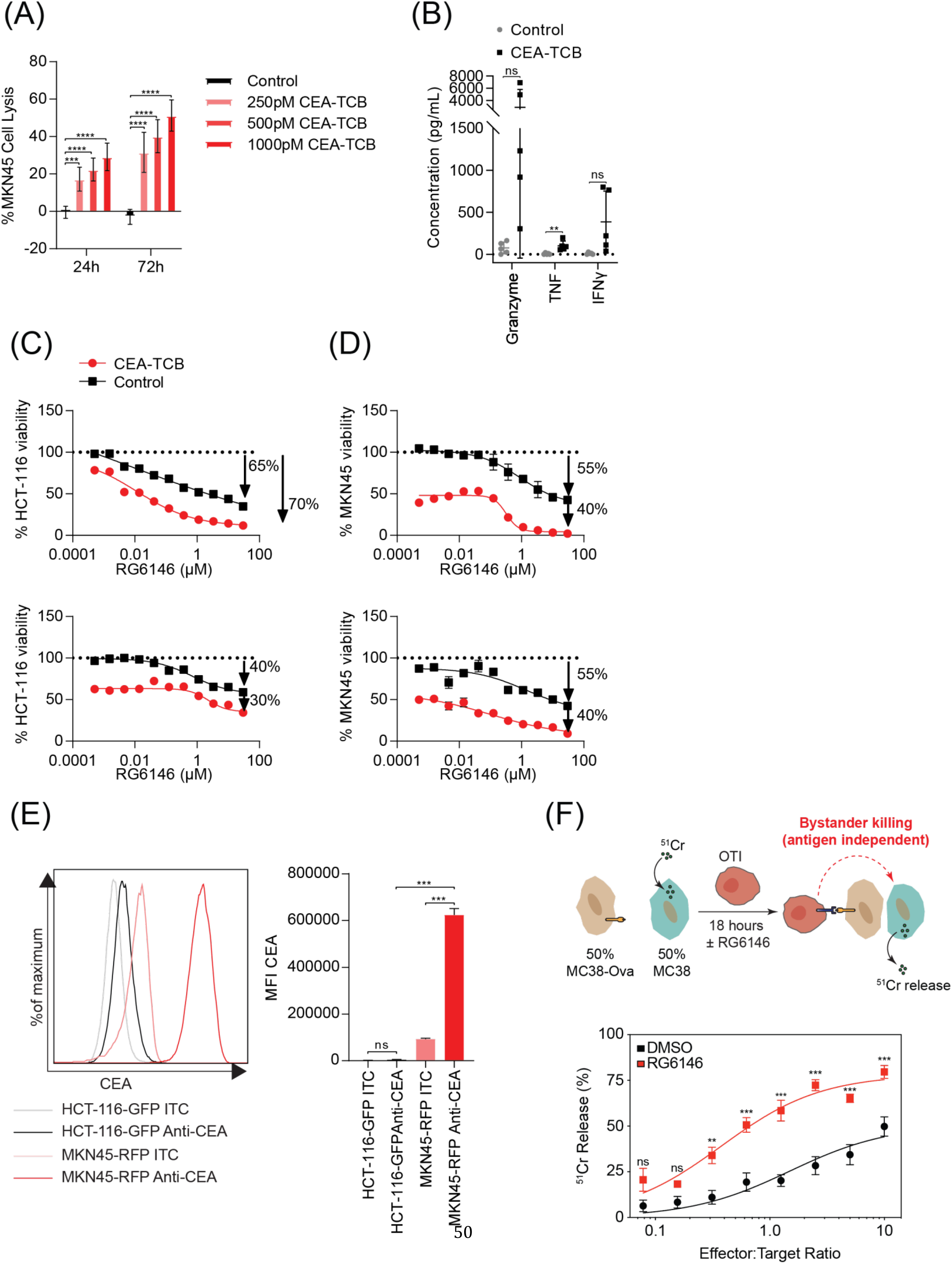
BET inhibition promotes antigen-independent by-stander killing. **(A)** Co-culture of MKN45 cells and PBMCs treated with 250-1000pM CEA-TCB, or vehicle control. Cell lysis of MKN45 cells (%) was assessed by LDH release into the supernatant after 24 and 72 hours of treatment. Data represents mean of three independent experiments. Statistical significance denotes difference between CEA-TCB and control (2-way ANOVA, ***p<0.001, ****p<0.0001). **(B)** Flow cytometry measurement of Granzyme B, TNF and IFNy concentration (pg/mL) in the supernatant of a co-culture assay with MKN45 cells and PBMCs treated for 24 hours with 20nM CEA-TCB or vehicle control. Mean of five independent experiments is shown. Statistical significance denotes difference between CEA-TCB and control treated cells (Unpaired T-Test, **p<0.01). **(C-D)** Viability of HCT-116 (C) and MKN45 (D) cells following treatment with the supernatant from a co-culture assay in the presence or absence of 20nM CEA-TCB and increasing concentrations of RG6146. Data was normalized to DMSO control and shows two independent PBMC Donors. **(E)** Flow cytometry analysis of CEA expression on HCT-116-GFP and MKN45-RFP cells. Mean of three independent experiments is shown. Statistical significance denotes difference between CEA antibody and ITC as well as CEA level between cell lines (2-way ANOVA, ***p<0.001). **(F)** Schematic representation of MC38 bystander killing assays where MC38-Ova cells are mixed 1:1 with MC38 parental cells labelled with chromium (^51^Cr) prior to co-culture with OTI T-cells in the presence or absence of RG6146 (2.5µM) for 18 hours. Release of ^51^Cr into the co-culture supernatant is then detected by liquid scintillation. Data represents three independent experiments. Statistical significance denotes difference between RG6146 and DMSO at each E:T ratio (2-way ANOVA, **p<0.01, ***p<0.001).

**Supplementary figure 8.**
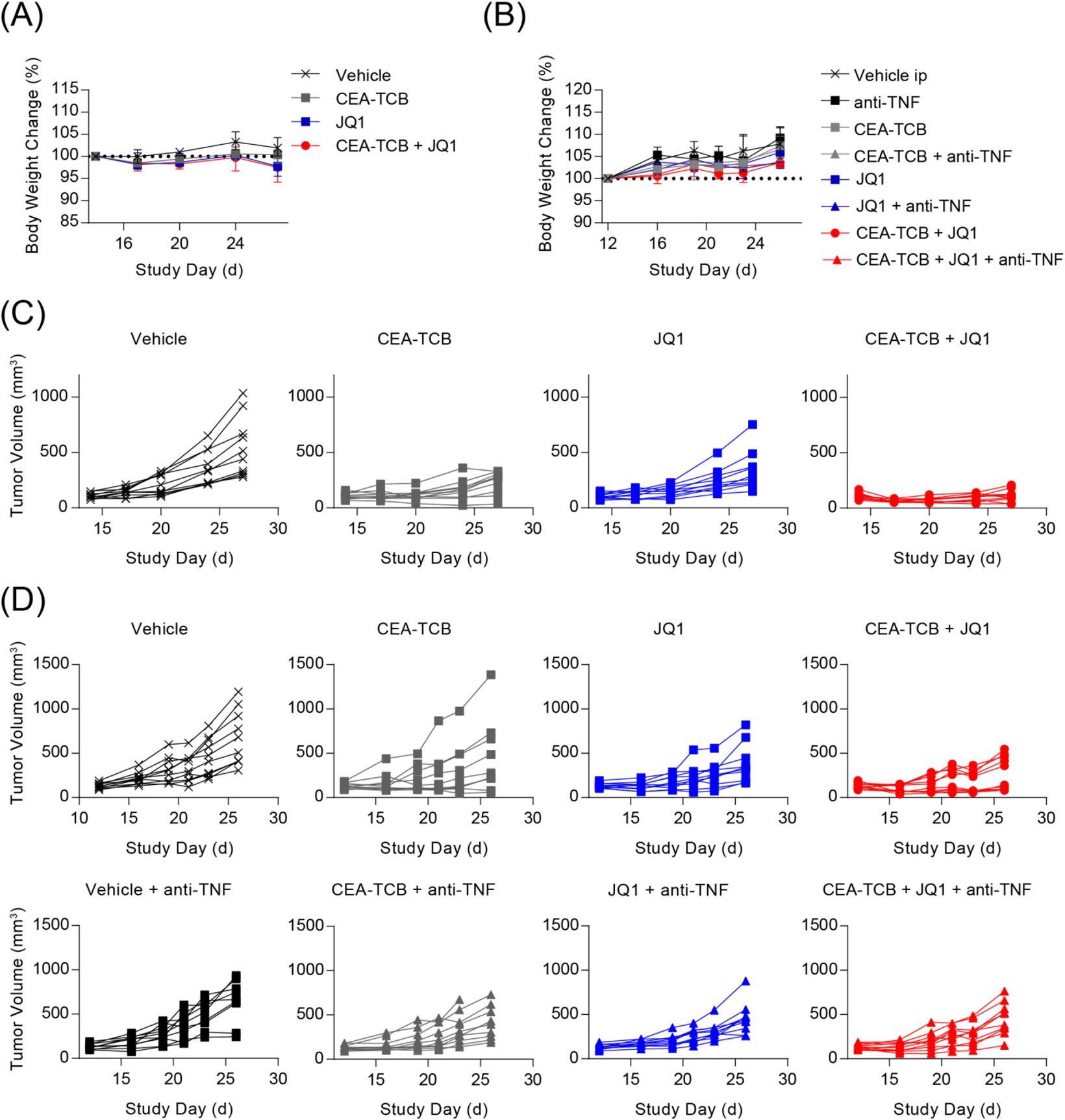
The activity of BET inhibitors and TCB antibodies is dependent on TNF. **(A)** Cohorts of transgenic C57BL/6 mice expressing human CEA (huCEA) mice (n=10 per treatment group) were transplanted with 5×10^5^ MC38 cells expressing CEACAM5. When tumor size reached 100-130 mm^3^, treatment commenced with 2.5mg/kg CEA-TCB antibody twice weekly, 50mg/kg JQ1 once daily, or the combination of CEA-TCB and JQ1. Change in body weight (%) was monitored during the course of the study. **(B)** Body weight monitoring from an experiment described in *A* with the addition of a TNF neutralizing antibody (anti-TNF) twice weekly. **(C)** Tumor growth curves from individual mice in the experiment described in *A*. **(D)** Tumor growth curves from individual mice in the experiment described in *B*

**Supplementary figure 9.**
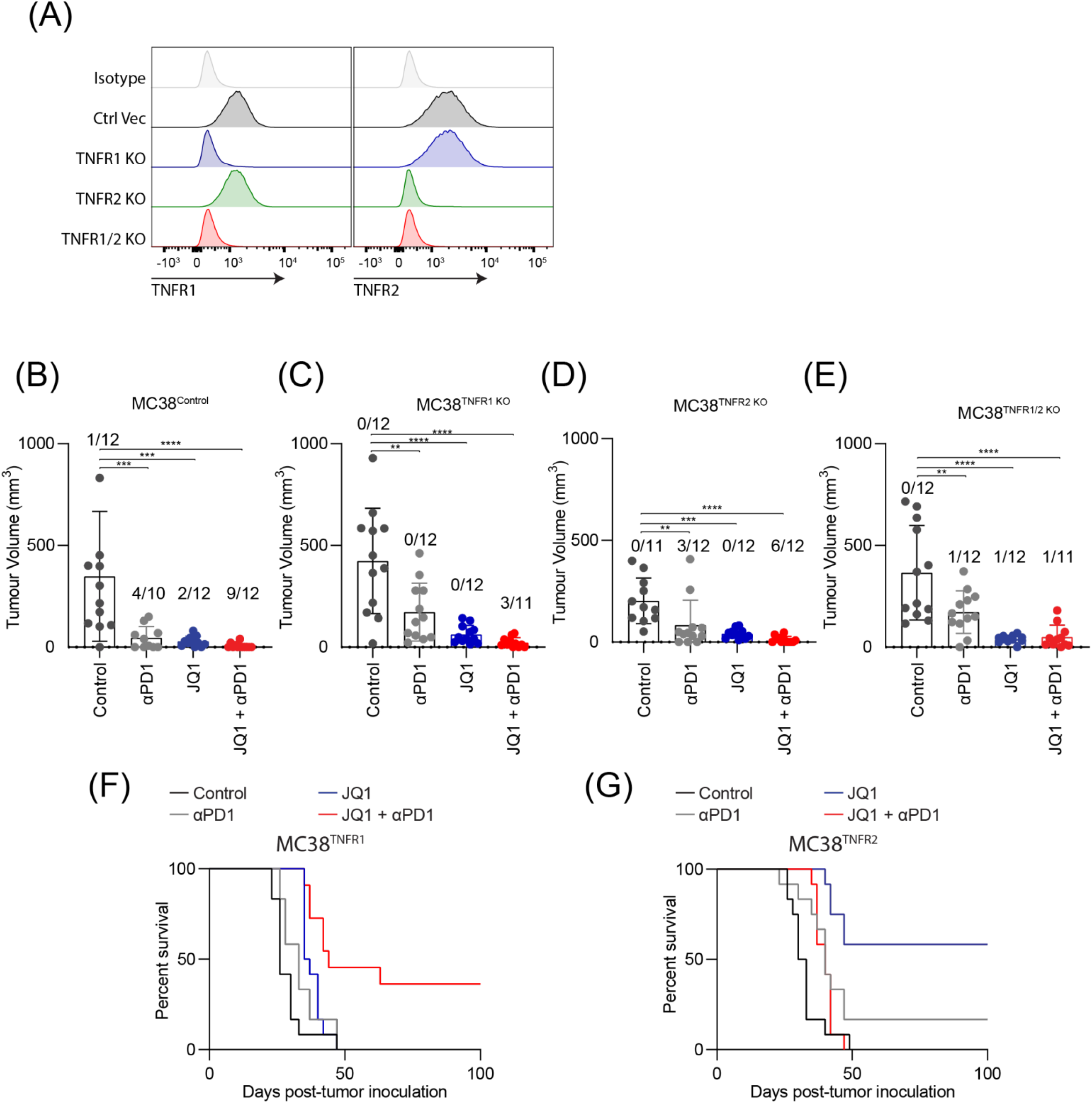
The activity of BET inhibitors and PD1 antibodies is dependent on TNF. (**A)** Histograms of MC38 cells depleted of TNFR1 (TNFR1 KO), TNFR2 (TNFR2 KO), both (TNFR1/2 KO) or vehicle control (Ctrl Vec) by electroporation of Cas9-sgRNA ribonucleoprotein complexes and stained for either TNFR1 or TNFR2 by flow cytometry. (B-E) Wild-type C57BL/6 mice bearing MC38 tumors depleted of (B) Control (C) TNFR1 (MC38 TNFR1 KO) (D) TNFR2 (MC38 TNFR2 KO) or (E) TNFR1 and TNFR2 (MC38_TNFR1/2 KO) were treated daily with 50mg/kg JQ1 or DMSO control alone (i.p.) and in combination with 0.2mg/kg anti-PD1 (i.p.) twice weekly. Bar chart showing tumor volume at day 17 of treatment. Data represents mean of two independent experiments (+/- SEM). Statistical significance denotes difference between treatment groups (1-way ANOVA, **p<0.01, ***p<0.001, ****p<0.0001). Numbers indicate mice that have cleared the tumor within each group. **(F-G)** Kaplan-meier survival curve of experiment described in *B* showing the overall survival of mice. Data represents mean of two independent experiments.

